# Single-Cell Signalling Analysis of Heterocellular Organoids

**DOI:** 10.1101/659896

**Authors:** Xiao Qin, Jahangir Sufi, Petra Vlckova, Pelagia Kyriakidou, Sophie E. Acton, Vivian S. W. Li, Mark Nitz, Christopher J. Tape

## Abstract

Organoids are powerful biomimetic tissue models. Despite their widespread adoption, methods to analyse cell-type specific post-translational modification (PTM) signalling networks in organoids are absent. Here we report multivariate single-cell analysis of cell-type specific signalling networks in organoids and organoid co-cultures. Simultaneous measurement of 28 PTMs in >1 million single small intestinal organoid cells by mass cytometry reveals cell-type and cell-state specific signalling networks in stem, Paneth, enteroendocrine, tuft, goblet cells, and enterocytes. Integrating single-cell PTM analysis with Thiol-reactive Organoid Barcoding *in situ* (TOB*is*) enables high-throughput comparison of signalling networks between organoid cultures. Multivariate cell-type specific PTM analysis of colorectal cancer tumour microenvironment organoids reveals that *shApc*, *Kras*^*G12D*^, and *Trp53*^*R172H*^ cell-autonomously mimic signalling states normally induced by stromal fibroblasts and macrophages. These results demonstrate how standard mass cytometry workflows can be modified to perform high-throughput multivariate cell-type specific signalling analysis of healthy and cancerous organoids.

## INTRODUCTION

Organoids are self-organising 3D tissue models comprising stem and differentiated cells (1). Organoid monocultures typically contain one major cell class (e.g. epithelial) and can be co-cultured with heterotypic cell-types (e.g. mesenchymal (2) or immune cells (3)) to model cell-cell interactions *in vitro*. When compared with traditional 2D cell culture, organoids more accurately represent their parental tissue and are emerging as powerful models for studying multicellular diseases such as cancer (4).

Post-translational modification (PTM) signalling networks underpin fundamental biological phenotypes and are frequently dysregulated in disease (5). As different cell-types have different signalling networks (6, 7), organoids likely contain several cell-type specific PTM networks that are essential to their biology. In order to fully utilise biomimetic models of healthy and diseased tissue, we must be able to study PTM signalling networks within organoids. Unfortunately, no technology currently exists to analyse cell-type specific PTM networks in organoids and organoid cocultures.

Organoids present several technical challenges over traditional 2D cultures for PTM analysis. Firstly, organoids are embedded in a protein-rich extracellular matrix (ECM) that confounds the application of phosphoproteomic analysis by liquid chromatography tandem mass spectrometry (LC-MS/MS). Organoids can be removed from ECM prior to LC-MS/MS, but as dissociation of live cells alters cell signalling (8), PTM measurements from dissociated live organoids do not truly represent *in situ* cellular states. Ideally, organoids should be fixed *in situ* to preserve PTM signalling, but LC-MS/MS analysis of heavily cross-linked phosphoproteomes is extremely challenging. Secondly, as organoids comprise multiple cell-types (e.g. stem and differentiated) and cell-states (e.g. proliferating, quiescent, and apoptotic), bulk phosphoproteomics cannot capture the biological heterogeneity present in organoids and organoid co-cultures (9). Although single-cell RNA-sequencing (scRNA-seq) can describe organoid cell-types (10), it cannot measure intracellular PTM signalling at the protein level. Finally, as signalling networks comprise multiple PTM nodes, low-dimensional methods (e.g. fluorescent imaging) cannot capture the complexity of PTM signalling networks (9). Collectively, to study PTM networks in organoids, we require signalling data that is: 1) cell-type specific, 2) derived from cells fixed *in situ*, and 3) measures multiple PTMs simultaneously.

Mass cytometry (MC, also known as cytometry time-of-flight (CyTOF)) uses heavy metal-conjugated antibodies to measure >35 proteins in single cells (11). Although MC is traditionally used for high-dimensional immunophenotyping, MC can also measure PTMs in heterocellular systems (e.g. peripheral blood mononuclear cells (PBMCs) (12) and tissue (8)). Given MC’s capacity to measure PTMs in mixtures of fixed cells, we theorised that MC workflows typically applied to immunophenotyping could be modified to study cell-type specific signalling networks in organoids.

Here we report the development of a custom multivariate-barcoded MC method to measure single-cell signalling in epithelial organoids and organoids co-cultured with stromal and immune cells. This method reveals that intestinal organoids display cell-type specific signalling networks that are intimately linked with cell-state. When applied to colorectal cancer (CRC) tumour microenvironment (TME) organoid co-cultures, we discovered that epithelial oncogenic mutations mimic signalling networks normally induced by stromal cells. These results demonstrate how a modified MC method can enable powerful multivariate single-cell analysis of cell-type specific signalling in heterocellular organoids.

## RESULTS

### Single-Cell Analysis of Organoids by Mass Cytometry

No technology currently exists to study cell-type specific protein signalling networks in organoids. Given its capacity to measure multiple PTMs in single cells, we hypothesised MC could be modified to study cell-type and cell-state specific signalling in organoids. To test this, we first developed a MC platform to measure single-cell signalling in the classical small intestinal organoid (13).

In this method, we first pulse live organoids with ^127^5-Iodo-2’-deoxyuridine (^127^IdU) to identify S-phase cells (14), fix organoids in Matrigel to preserve cell signalling, and stain organoids with ^194/8^Cisplatin to label dead epithelia (15). Using a custom workflow, we then physically and enzymatically dissociate the fixed organoids into single cells prior to extra- and intracellular heavy-metal antibody staining (Fig. 1a). We next performed a comprehensive screen for intestinal epithelial cell-type identification antibodies including stem (LGR5, LRIG1, OLFM4), Paneth (Lysozyme), goblet (MUC2, CLCA1), enteroendocrine (CHGA, Synaptophysin), tuft cells (DCAMKL1), and enterocytes (FABP1, Na/K-ATPase) that bind fixed antigens and are compatible with rare-earth metal conjugation for MC. Cell-type identification antibodies were validated by organoid directed differentiation (16) (Supplementary Fig. 1) and integrated into a panel of 28 anti-PTM rare-earth metal antibodies spanning multiple core signalling nodes (Supplementary Table 1, 45 parameters (40 antibodies) / cell). When analysed by MC, this method enables the measurement of 28 signalling PTMs across 6 cell-types in >1 million single cells from fixed intestinal organoids (Figs. 1b, c, 2a, and Supplementary Fig. 2a).

**Fig. 1.**
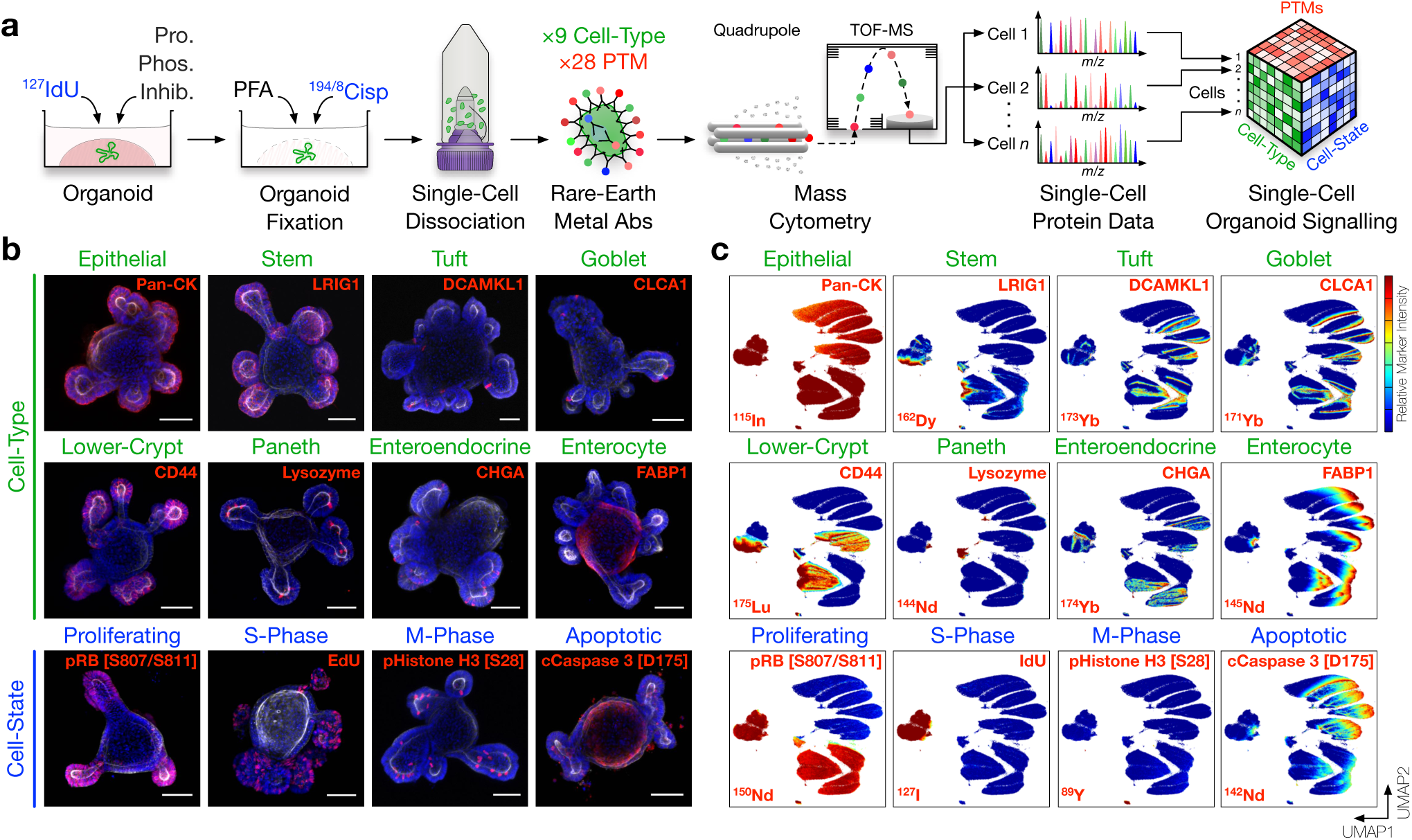
Cell-Type and Cell-State Identification of Single Organoid Cells by Mass Cytometry. **a)** Experimental workflow. Live organoids are pulsed with ^127^IdU to label S-phase cells, treated with protease / phosphatase inhibitors, fixed with PFA to preserve post translational modification (PTM) signals, and stained with ^194/8^Cisplatin to label dead cells. Fixed organoids are then dissociated into single cells, stained with rare-earth metal-conjugated antibodies, and analysed by single-cell mass cytometry (MC). The resulting dataset contains integrated cell-type, cell-state, and PTM signalling information. **b)** Confocal immunofluorescence (IF) of small intestinal organoids stained with rare-earth metal-conjugated MC antibodies highlighting individual cell-type and cell-state markers (red), F-Actin (white), and DAPI (blue), scale bars = 50 µm. (See Supplementary Fig. 1 for antibody validation via directed differentiation.) **c)** UMAP (Uniform Manifold Approximation and Projection) distribution of 1 million single organoid cells analysed by MC resolves six major intestinal cell-types across proliferating, S-phase, M-phase, and apoptotic cell-states. Colours represent normalised local parameter intensity. (See Supplementary Fig. 2 for cell-type and cell-state classification.)

**Fig. 2.**
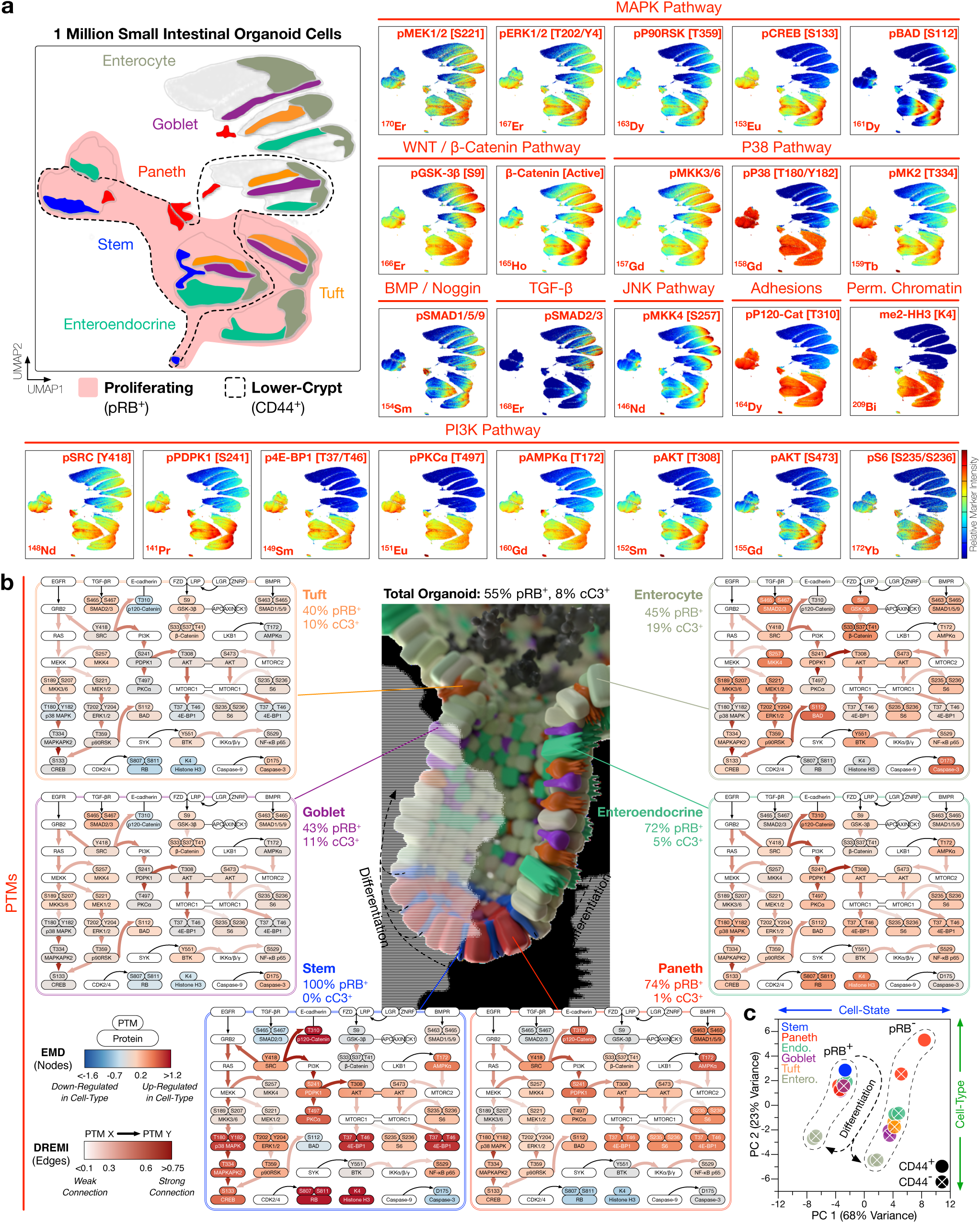
Cell-Type and Cell-State Specific Signalling Analysis of Intestinal Organoids. **a)** UMAP distributions of PTMs across 1 million single organoid cells analysed by MC. Cell-type and cell-state UMAP guide is shown top left (see Supplementary Fig. 2 for cell-type and cell-state classification). Combining cell-type, cell-state, and PTM measurements enables cell-type specific analysis of intestinal organoid signalling. Colours represent normalised local parameter intensity. **b)** Cell-type specific PTM signalling networks in small intestinal organoids, with nodes coloured by PTM-EMD (Earth Mover’s Distance) scores quantifying PTM intensity (relative to all organoid cells) and edges coloured by DREMI (Density Resampled Estimation of Mutual Information) scores quantifying PTM-PTM connectivity. Small intestinal organoids display cell-type specific signalling networks. **c)** Principal Component Analysis (PCA) of PTM-EMDs for all organoid cell-types, either proliferating (pRB^+^) / G0 (pRB^−^) or in lower-crypts (CD44^+^) / villi (CD44^−^). Organoid cell signalling is dictated by cell-state (PC 1) and cell-type (PC 2). (See Supplementary Fig. 3 for complete EMD-DREMI signalling maps.)

Combining ^194/8^Cisplatin and ^127^IdU with canonical cell-cycle markers (e.g. pRB [S807/S811], Cyclin B1, and pHistone H3 [S28] (14)) allows clear identification of live / dead cells and classification of single organoid cells into cell-cycle stages including G0, G1, S, G2, and M-phase (17) (Supplementary Fig. 2b). Integrated cell-type and cell-state data from small intestinal organoids confirmed that stem and Paneth cells are largely proliferative (pRB^+^, cCaspase3 [D175]^−^), whereas differentiated epithelia are often post-mitotic (pRB^−^, IdU^−^ / pHH3^−^) or apoptotic (cCaspase3^+^) (Fig. 1c). Consistent with the finding that intestinal progenitor cells have permissive chromatin *in vivo* (18), proliferating intestinal organoid cells also present H3K4me2 whereas post-mitotic cells do not (Fig. 2a). These results confirmed that a modified MC workflow can provide cell-type and cell-state specific information from millions of single organoid cells.

### Cell-Type and Cell-State Specific Signalling Networks in Intestinal Organoids

Following cell-type and cell-state identification, we next sought to construct cell-type specific PTM signalling networks in small intestinal organoids. To investigate whether stem, Paneth, enteroendocrine, tuft, goblet cells, and enterocytes employ different PTM signalling networks, we combined Earth Mover’s Distance (EMD) (19, 20) and Density Resampled Estimation of Mutual Information (DREMI) (21) to build quantitative cell-type specific signalling networks from single-cell organoid PTM data (Fig. 2b and Supplementary Fig. 2). In these networks, EMD quantifies PTM intensity (node score) for each organoid cell-type relative to the total organoid population and DREMI quantifies PTM-PTM connectivity (edge score) within the network.

EMD-DREMI analysis revealed cell-type specific PTM signalling networks in small intestinal organoids. As canonical WNT signalling is mainly driven by protein interactions, localisation, and degradation (22) – not a classical PTM cascade – MC is not well suited to studying the WNT pathway. Despite this limitation, evidence of WNT flux via inhibited pGSK-3*β* [S9] and non-phosphorylated *β*-Catenin is observed in all organoid cell-types (Fig. 2a). In contrast, MAPK and PI3K pathways display unexpected cell-type specificity. For example, stem cells channel MAPK signalling through pERK1/2 [T202/Y204], pP90RSK [T359], and pCREB [S133], but fail to connect with pBAD [S112] (Fig. 2a, b). On the contrary, differentiated epithelia direct MAPK signalling away from pCREB and towards pBAD when proliferating, and lose all MAPK activity in G0 and apoptosis (Fig. 2a). Despite their strong mitogenic signalling profile, stem cells are unique among proliferating cells in their failure to phosphorylate BAD. This suggests that intestinal stem cells avoid apoptosis independent of the classical BAD-BCL-BAX/BAK axis and may compensate via high MAPK and P38 flux to CREB. Consistent with the observation that PI3K signalling is important for intestinal crypt cells (23), stem and Paneth cells are enriched for pSRC [Y418] and downstream PI3K effectors such as pPDPK1 [S241], pPKC*α* [T497], pAKT [T308] / [S473], and p4E-BP1 [T37/T46] (Fig. 2a, b). Despite the presence of the BMP inhibitor Noggin in organoid culture media, Paneth cells display unexpectedly high BMP signalling (via pSMAD1/5 [S463/S465] and SMAD9 [S465/S467]) (Fig. 2a, b). This observation suggests that Paneth cells are either hypersensitive to BMP ligands or can cell-intrinsically activate SMAD1/5/9 (possibly via inhibition of SMAD phosphatases).

Several PTM signalling events correlate with cell-state in intestinal organoids. For example, irrespective of cell-type or location, pP38 MAPK [T180/Y182] and pP120-Catenin [T310] are active in all proliferating cells, and both pAKT [T308] and pMKK4 [S257] are hyperactivated in M-phase (Figs. 1c and 2a). In contrast, TGF-*β* signalling (via pSMAD2 [S465/S467] and SMAD3 [S423/S425]) is exclusively active in post-mitotic epithelia (Fig. 2a), consistent with TGF-*β*’s role in epithelial growth-arrest (24). To investigate the relationship between cell-type and cell-state in PTM signalling networks, we performed principal component analysis (PCA) of PTM-EMDs for each organoid cell-type, either proliferating or in G0 (pRB^+/–^), located in lower-crypts or villi (CD44^+/–^). PCA revealed that both cell-state (PC1, 68% variance) and, to a lesser extent, cell-type (PC2, 23% variance) dictate cell-signalling in small intestinal organoids (Fig. 2c and Supplementary Fig. 3). This analysis demonstrates that both cell-type and cell-state are intimately linked with cell-signalling and warns against bulk PTM analysis of organoids where cell-type and cell-state resolution is lost. Collectively, these results confirmed that MC can identify novel cell-type and cell-state specific signalling networks in small intestinal organoids and underscore the importance of single-cell data when studying heterogenous systems such as organoids.

### Single-Cell Organoid Multiplexing using Thiol-reactive Organoid Barcoding *in situ* (TOB*is*)

We have demonstrated how a modified MC platform can be applied to cell-type and cell-state specific signalling measurement in organoids. However, in order to study differential signal transduction in organoid models of healthy and diseased tissue, we must also be able to directly compare PTM networks between different organoid cultures. In addition to high dimensional single-cell PTM measurements, a major advantage of MC is its ability to perform multiplexed barcoding of experimental variables (25, 26). Unfortunately, commercially available Palladium-based barcodes cannot bind organoids *in situ* (Supplementary Fig. 4a) as they react with Matrigel proteins (Supplementary Fig. 4b), meaning that organoids must be removed from Matrigel and dissociated separately before barcoding. Individually removing fixed organoids from Matrigel is a very low-throughput process that limits the scalability of organoid MC multiplexing. We theorised that if organoids could be barcoded *in situ*, barcoded organoids could be pooled, dissociated, and processed as a single high-throughput MC sample. To explore this idea, we developed a new strategy to isotopically barcode organoids while still in Matrigel.

MC barcoding strategies can use amine-(26) or thiol-reactive (25) chemistries. We first used fluorescent probes to investigate how each of these chemistries react with ECM proteins and organoids. Amine-reactive probes (Alexa Fluor 647 NHS ester) bind ECM proteins (via lysines and N-terminal amines) and thus fail to label organoids in Matrigel. In contrast, thiol-reactive probes (Alexa Fluor 647 C_2_ maleimide) bypass ECM proteins and bind exclusively to reduced-cysteines on organoids *in situ* (Fig. 3a, Supplementary Fig. 4c). We subsequently confirmed that thiol-reactive monoisotopic mass-tagged probes (C_2_ maleimide-DOTA-^157^Gd) also bind organoids *in situ*, whereas amine-reactive probes (NHS ester-DOTA-^157^Gd) only react *ex situ* (Fig. 3b). This data confirmed that thiol-reactive chemistries can be used to barcode organoids while still in Matrigel (Fig. 3c). Using this knowledge, we developed a 20-plex (*6*-choose-*3*, doublet-filtering (25, 26)) thiol-reactive barcoding strategy based on monoisotopic tellurium maleimide (TeMal) (27) (^124^Te, ^126^Te, ^128^Te, ^130^Te) and Cisplatin (28) (^196^Pt, ^198^Pt) that can bypass Matrigel proteins and bind directly to fixed organoids *in situ* (Fig. 3d and Supplementary Fig. 4d). This <u>T</u>hiol-reactive <u>O</u>rganoid <u>B</u>arcoding *<u>i</u>n <u>s</u>itu* (TOB*is*) approach enables high-throughput multivariate single-cell organoid signalling analysis in a single tube (Fig. 3e).

**Fig. 3.**
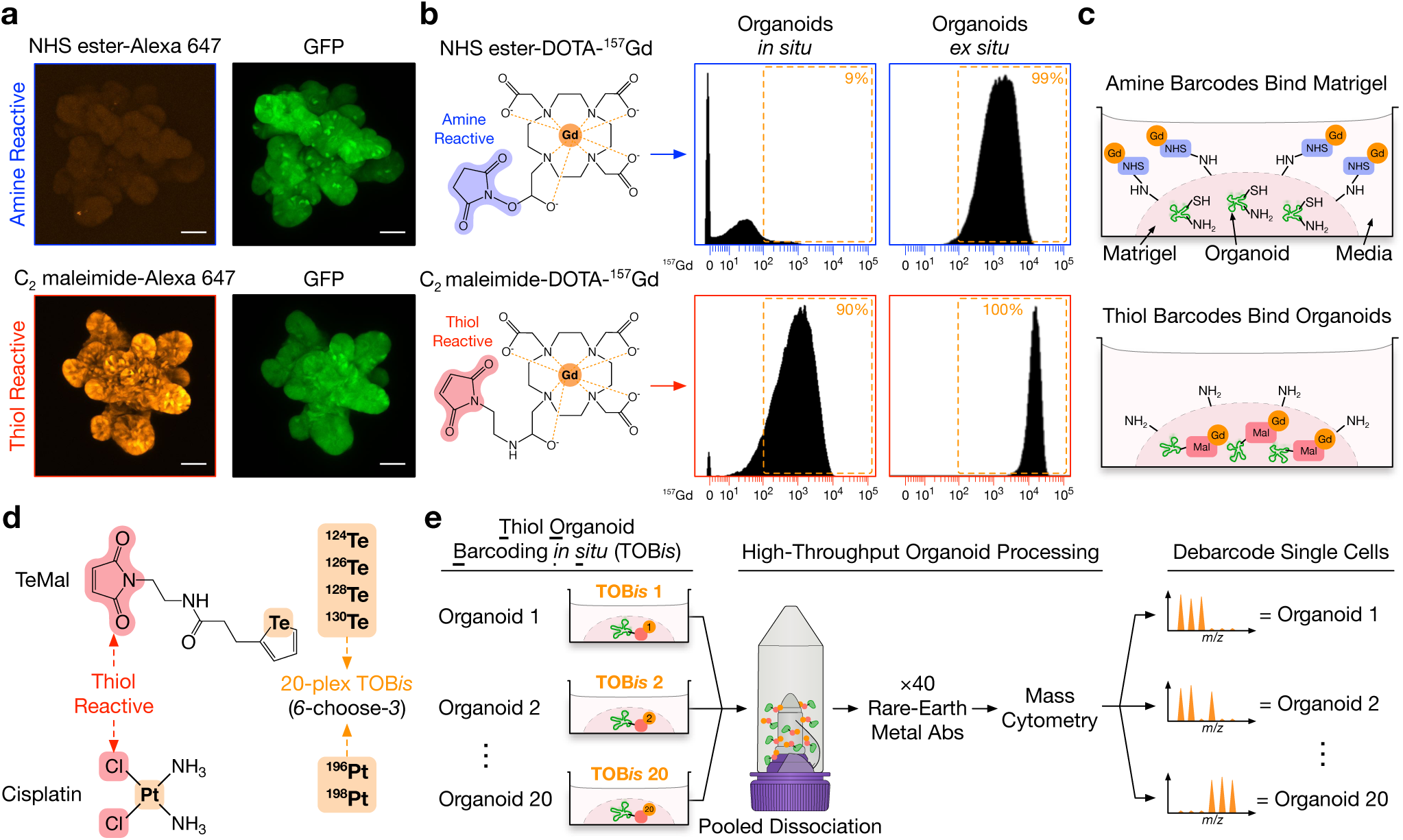
Thiol-Reactive Organoid Barcoding *in situ* (TOB*is*) for Single-Cell Organoid Multiplexing. **a)** Confocal IF of fixed GFP^+^ small intestinal organoids stained with either amine-reactive fluorescent probe Alexa Fluor 647 NHS ester or thiol-reactive fluorescent probe Alexa Fluor 647 C_2_ maleimide while still in Matrigel, scale bars = 50 µm. Amine-reactive probes bind diffusely to Matrigel with very poor binding to organoids, whereas thiol-reactive probes bypass Matrigel and bind directly to organoids (see Supplementary Fig. 4c for Matrigel background staining). **b)** Small intestinal organoids stained with either amine-reactive NHS ester-DOTA-157Gd or thiol-reactive C_2_ maleimide-DOTA-157Gd *in situ* (still in Matrigel) or *ex situ* (removed from Matrigel) and analysed by MC. While both probes bind organoid cells *ex situ*, only thiol-reactive C_2_ maleimide-DOTA-^157^Gd bind organoids *in situ*. **c)** Model of amine- and thiol-reactive barcodes in organoid culture. **d)** Thiol-reactive tellurium maleimide (TeMal) (^124^Te, ^126^Te, ^128^ Te, ^130^Te) and Cisplatin (^196^Pt, ^198^Pt) isotopologs combined to form a 20-plex (*6*-choose-*3*) doublet-filtering barcoding strategy. **e)** Thiol-Reactive Organoid Barcoding *in situ* (TOB*is*) workflow. When combined with cell-type, cell-state, and PTM probes, TOB*is* allows organoids to be barcoded while still in Matrigel and rapidly processed as a single sample. (See Supplementary Fig. 5 for additional details.)

It is worth noting that as Te and Pt metals are not typically conjugated to antibodies in MC, TOB*is* multiplexing does not compromise the number of antigens being measured. Moreover, unlike commercially available Pd barcodes (occupying ^102^Pd, ^104^Pd, ^105^Pd, ^106^Pd, ^108^Pd, and ^110^Pd channels), TOB*is* barcodes do not clash with the recently developed Cadmium antibody labelling metals (^106^Cd, ^110^Cd, ^111^Cd, ^112^Cd, ^113^Cd, ^114^Cd, and ^116^Cd). Furthermore, as barcoding is performed on fixed organoids embedded within Matrigel, TOB*is* does not require the numerous centrifugation or cell membrane permeabilisation steps used in traditional solution-phase barcoding. This greatly increases organoid sample-throughput (Supplementary Fig. 5a–d) and single-cell recovery (Supplementary Fig. 5e–g), thereby facilitating high-throughput organoid MC applications.

### Multivariate Cell-Type Specific Signalling Analysis of Intestinal Organoid Development

Traditional mass-tag barcoding allows direct comparison of solution-phase cells (e.g. immune cells) between experimental conditions (26). When combined with cell-type, cell-state, and PTM probes, TOB*is* multiplexing now enables PTM signalling networks to be directly compared between organoid cultures in a high-throughput manner. To demonstrate this, we applied TOB*is* to study cell-type specific epithelial signalling during 7 days of small intestinal organoid development (Fig. 4 and Supplementary Table 1, 50 parameters (40 antibodies) / cell).

**Fig. 4.**
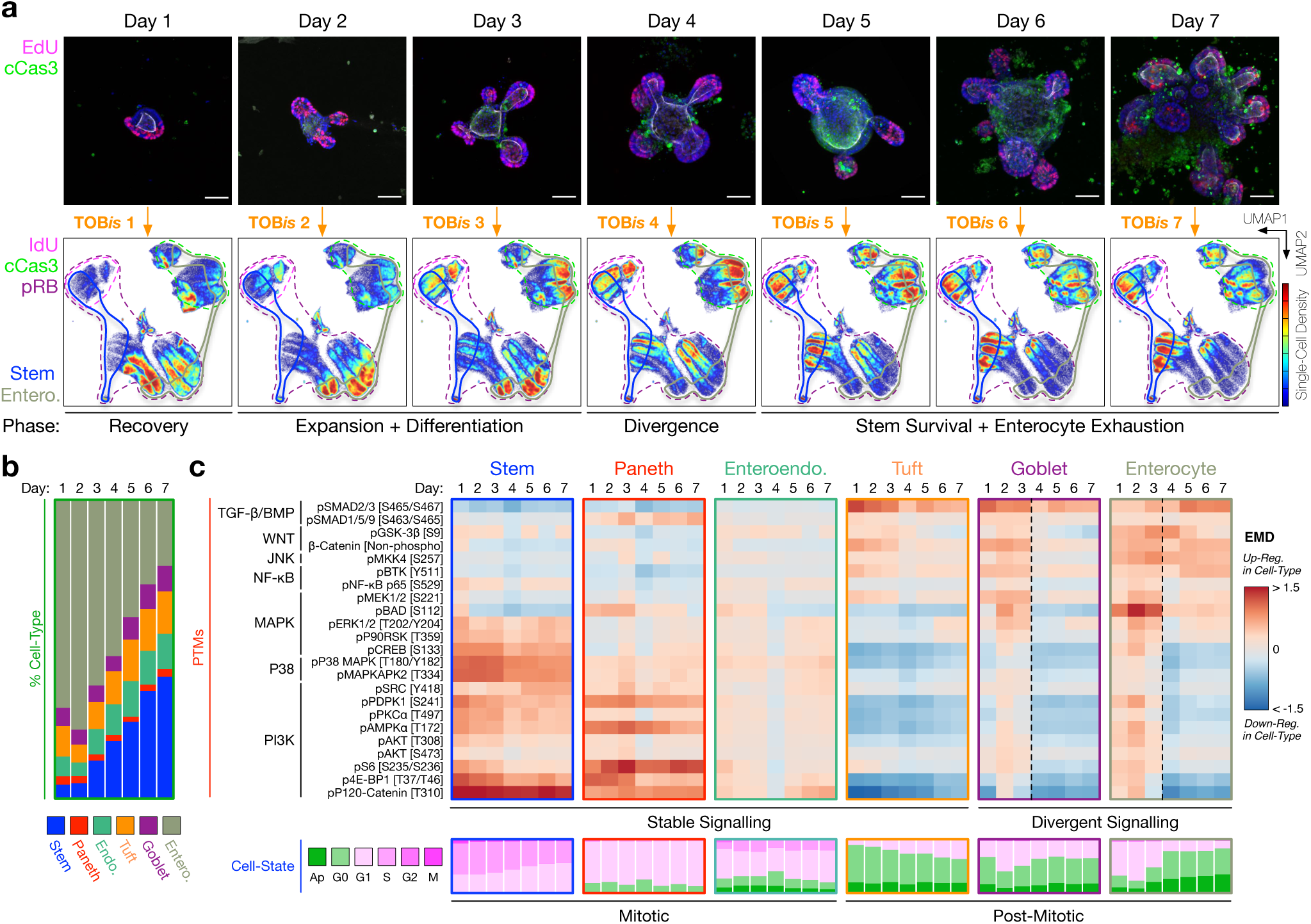
Cell-Type Specific Signalling During Intestinal Organoid Development. **a)** Time-course confocal IF of intestinal organoid development illustrating S-phase (EdU^+^, magenta) and apoptotic (cCaspase 3 [D175]^+^, green) cells, scale bars = 50 µm. Each time point was barcoded by TOB*is*, pooled into a single sample, and analysed by MC. Cell-density UMAP distributions of 2 million single organoid cells reveal changes in cell-type and cell-state during organoid development. **b)** Cell-type composition of small intestinal organoids during development. Stem cells accumulate at the expense of enterocytes during organoid culture. **c)** Cell-type specific PTMs and cell-states of stem, Paneth, enteroendocrine, tuft, goblet cells, and enterocytes during intestinal organoid development. Cell-state analysis shows the proportions of apoptotic, G0-, G1-, S-, G2-, and M-phase cells. Irrespective of time point, stem, Paneth, and enteroendocrine cells are stably mitotic, whereas tuft, goblet cells, and enterocytes are frequently post-mitotic. Stem, Paneth, enteroendocrine, and tuft cells display stable signalling over time, whereas goblet cell- and enterocyte-signalling diverge from Day 4.

Analysis of 28 PTMs from 2 million single organoid cells revealed that after 1 day of culture, organoids seeded as single crypts exist in a ‘recovery’ phase where 70% cells have entered the cell-cycle (pRB^+^), but <5% reach S-phase (IdU^+^) (compared to 20% in developed organoids) (Fig. 4a). Days 2 and 3 mark a rapid ‘expansion and differentiation’ phase of organoid development where stem, Paneth, goblet cells, and enterocytes activate MAPK, P38, and PI3K pathways – although stem cells again fail to inhibit BAD (Fig. 4c). By Day 4, intestinal organoids reach a critical ‘divergence’ phase where crypt and villus signalling digress dramatically. While stem and Paneth cells maintain active MAPK, P38, and PI3K pathways, enterocytes lose major PI3K (pPDPK1, p4E-BP1, pS6 [S235/S236], pAMPK*–* [T172], pSRC, and pPKC*–*) and P38 (pP38 MAPK, pMAPKAPK2 [T334], and pCREB) activity (Fig. 4c). As a result, by Days 5 to 7, enterocytes are largely post-mitotic or apoptotic (pRB^−^ / cCaspase3^+^), with high TGF-*β* signalling, whereas stem cells retain mitogenic flux and cell-cycle activity (Fig. 4c). Consequently, stem cell number increases while enterocytes become exhausted at the end of intestinal organoid development (Fig. 4a, b). Notably, both stem and Paneth cells continue to display high MAPK, P38, and PI3K activity even at this late stage of organoid culture (Fig. 4c). This suggests that maintaining a stable signalling flux is a core feature of intestinal crypt cells. In contrast, tuft cells display high TGF-*β* signalling, low MAPK/ P38 / PI3K activity, and low cell-cycle activity throughout organoid development (Fig. 4c). This implies that irrespective of organoid age, tuft cells shut down mitotic signalling pathways and terminally exit the cell cycle once differentiated. Such variations in organoid cell-state (Fig. 4a), cell-type (Fig. 4b), and PTM activity (Fig. 4c) suggest developmental stage should be carefully considered when performing organoid experiments. Collectively, this analysis revealed cell-type specific PTM signalling during intestinal organoid development and confirmed that TOB*is* can be used to perform multivariate single-cell signalling analysis of heterogenous organoid cultures.

### Single-Cell Signalling Analysis of CRC TME Organoids

We have demonstrated how a modified MC workflow enables high-throughput comparison of cell-type specific signalling networks in epithelial organoids. Given that MC can theoretically resolve any cell-type, we next expanded this platform to study PTM signalling in heterocellular organoid co-culture models of CRC.

CRC develops through successive oncogenic mutations – frequently resulting in loss of APC activity, activation of KRAS, and perturbation of TP53 (29). In addition to oncogenic mutations, stromal fibroblasts (30, 31) and macrophages (32) in the TME have also emerged as major drivers of CRC (33). While the underlying driver mutations of CRC have been well studied, how they dysregulate epithelial signalling relative to microenvironmental cues from stromal and immune cells is unclear.

To investigate this, we cultured wild-type (WT), *shApc* (A), *shApc* and *Kras^G12D/+^* (AK), or *shApc*, *Kras^G12D/+^*, and *Trp53^R172H/-^* (AKP) (34, 35) colonic epithelial organoids either alone, with colonic fibroblasts, and/or macrophages (Fig. 5a, b, and Supplementary Fig. 6). Each CRC genotype-microenvironment organoid culture was fixed, TOB*is*-barcoded, and single-cell signalling analysis was performed in one multivariate MC run (Fig. 5a and Supplementary Table 2, 50 parameters (40 antibodies) / cell). Addition of myeloid (CD68 and F4/80) and mesenchymal (Podoplanin) heavy-metal antibodies enabled clear resolution of epithelial organoids, macrophages, and fibroblasts from each barcoded condition (Fig. 5c). This experimental design allowed us to directly compare mutation- and microenvironment-driven cell-type specific signalling networks in CRC organoid mono- and co-cultures.

**Fig. 5.**
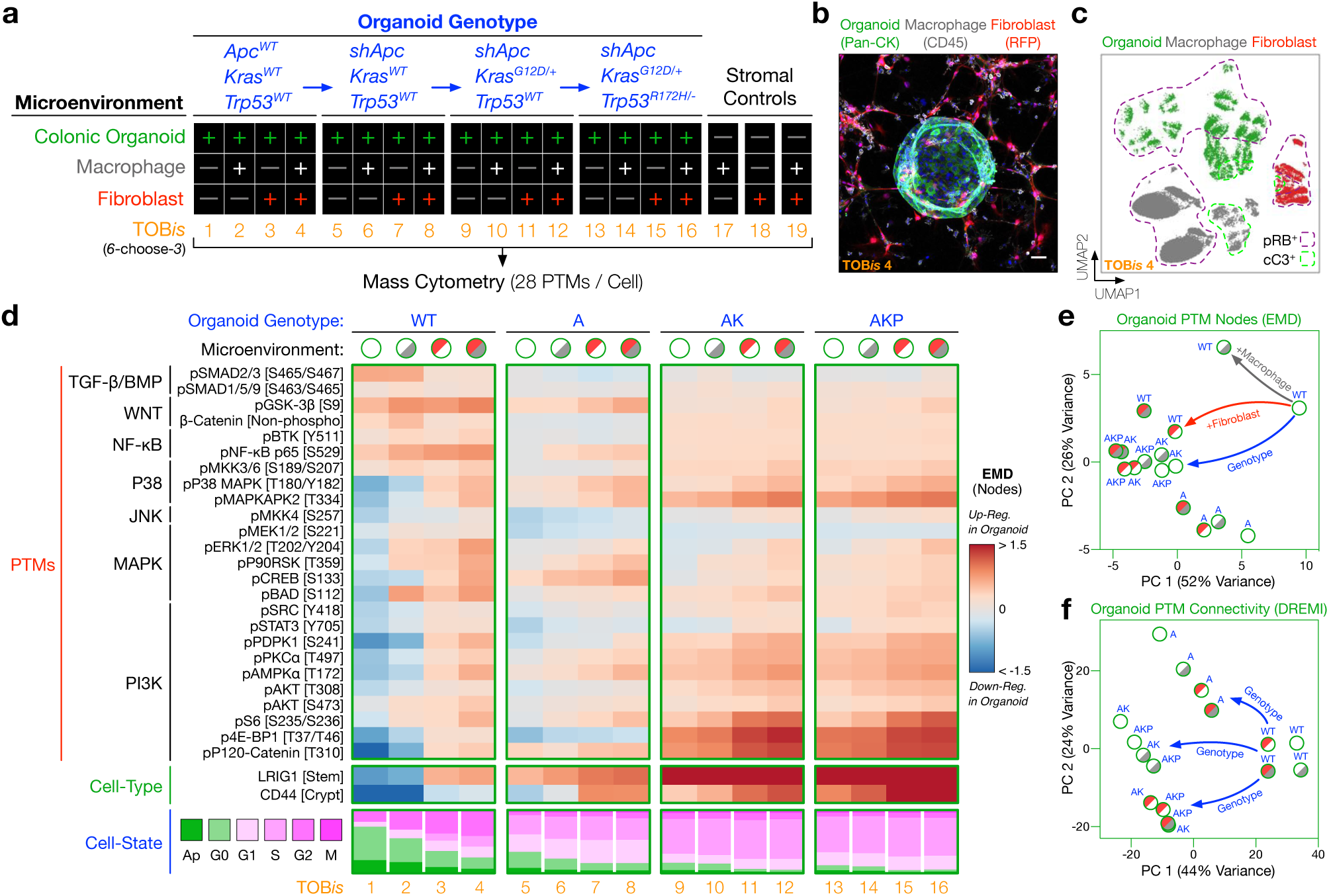
Single-Cell Signalling Analysis of Colorectal Cancer (CRC) Tumour Microenvironment Organoids. **a)** Experimental design. CRC organoid genotypes (wild-type (WT), *shApc* (A), *shApc* and *Kras*^*G12D/+*^ (AK), *shApc*, *Kras*^*G12D/+*^, and *Trp53*^*R172H/-*^(AKP)) were cultured in the presence or absence of colonic fibroblasts and/or macrophages (without exogenous growth factors). Each condition was TOB*is*-barcoded, pooled into a single sample, and analysed by MC (28 PTMs / cell). **b)** Confocal IF of a WT colonic organoid (Pan-CK, green) co-cultured with colonic fibroblasts (RFP, red), and macrophages (CD45, grey) (TOB*is* 4), scale bar = 50 µm. **c)** UMAP distribution of the colonic microenvironment model resolves single epithelial cells (green), fibroblasts (red), and macrophages (grey) (TOB*is* 4). **d)** PTMs, progenitor cell-types, and cell-states of colonic epithelial organoids across all genotype / microenvironment combinations. The grey and red shades in the microenvironmental conditions represent macrophages and fibroblasts respectively. (See Supplementary Figs. 7 and 8 for complete EMD-DREMI signalling maps of organoids, macrophages, and colonic fibroblasts.) **e)** PCA of 28 PTM-EMDs for colonic epithelial organoids across all genotype / microenvironment combinations. CRC organoids with AK / AKP mutations mimic the signalling flux driven by colonic fibroblasts. (See Supplementary Fig. 8c, d for PTM-EMD PCAs for macrophages and colonic fibroblasts.) **f)** PCA of 756 PTM-DREMIs for colonic epithelial organoids across all genotype / microenvironment combinations. Epithelial signalling connectivity is regulated by genotype rather than microenvironment. (See Supplementary Fig. 8e, f for PTM-DREMI PCAs for macrophages and colonic fibroblasts.)

As expected, oncogenic mutations have a large cell-autonomous effect on epithelial signalling. Although APC mutations are well known to upregulate WNT signalling (22), we found that the loss of APC also activates the P38 pathway (pP38 MAPK and pMAPKAPK2), downregulates TGF-*β* / BMP signalling (pSMAD2/3 and pSMAD1/5/9), and activates p120-Catenin in colonic organoids (Fig. 5d). Subsequent oncogenic *Kras*^*G12D/+*^ and *Trp53*^*R172H/-*^ mutations further cell-autonomously upregulate not only the classical MAPK pathway, but also major PI3K nodes (pPDPK1, pAKT, pS6, and p4E-BP1) (Fig. 5d). As a result, AK and AKP organoids display increased stem / progenitor cell-type markers LRIG1 and CD44, decreased apoptosis, and increased mitogenic cell-state relative to WT and A organoids (Fig. 5d).

Both oncogenes and stromal cells can dysregulate cancer cell signalling (7). However, to what extent this is driven by oncogenic mutations (cell-intrinsic) or the TME (cell-extrinsic) is less clear. To investigate this, we directly compared mutation- and microenvironment-driven signalling in CRC organoids. To our surprise, we found that microenvironmental cues have a comparable impact on PTM regulation to oncogenic mutations (Fig. 5e). In contrast, while mutations and stromal cells can both drive epithelial PTM activity, PTM-PTM connectivity is regulated largely by genotype, not microenvironment (Fig. 5f). This observation suggests that oncogenic mutations fundamentally re-wire signalling networks, whereas stromal cells regulate acute signalling flux. We also found that stromal cells further upregulate the PI3K pathway (pS6, p4E-BP1, and pAKT) in CRC organoids that already contain *Kras*^*G12D*^ and *Trp53*^*R172H*^ mutations (Fig. 5d and Supplementary Fig. 7). Microenvironmental hyper-activation of the epithelial PI3K pathway may contribute towards the poor prognosis of CRC patients with highly stromal tumours (30, 31).

In addition to mutation- and microenvironment-driven epithelial signalling, we discovered previously unreported polarity in fibroblast and macrophage cell-cell communication. For example, macrophage signalling pathways (MAPK, PI3K, and NF-*Ÿ*B) are heavily upregulated by fibroblasts (Supplementary Fig. 8a, c, e), whereas fibroblast signalling is scarcely altered by macrophages (Supplementary Fig. 8b, d, f). In contrast, epithelial cells upregulate MAPK and P38 signalling in fibroblasts, which in turn, reciprocally activate MAPK and P38 signalling in epithelial cells (Supplementary Fig. 7 and 8b). These results suggest that colonic fibroblasts are major regulators of intercellular signalling in the colonic microenvironment and should be further investigated as drivers of CRC.

### Oncogenic Mutations Mimic Stromal Signalling Networks

Cell-type specific PCA of EMD-PTMs suggested that mutation- and microenvironment-driven signalling in colonic organoids are related (Fig. 5e). To further investigate the parity between genotypic and microenvironmental regulation of epithelial signalling, we overlaid single-cell MC data from WT, A, AK, and AKP organoids onto a fixed-node microenvironmental Scaffold map (36) constructed from WT colonic organoids alone or co-cultured with colonic fibroblasts and/or macrophages (Fig. 6a and Supplementary Fig. 9a). This unsupervised analysis confirmed that *Apc*, *Kras*, and *Trp53* oncogenic mutations mimic the signalling profile of WT organoids in the presence of stromal cells. Inverted organoid genotype Scaffold maps also expose a striking similarity between mutation- and microenvironment-driven signalling (Supplementary Fig. 9b). Direct comparison of organoid PTMs revealed that both PI3K / PKC (pPDPK1, pPKC*–*, pAKT, p4E-BP1, pS6, pSRC, pP120-Catenin, and pAMPK*–*) and P38 / MAPK (pP38 MAPK, pMAPKAPK2, pP90RSK, pCREB, and pBAD) nodes are analogously up-regulated by oncogenic mutations and microenvironmental cues (Figs. 5d and 6b). Activation of these pathways by either oncogenic mutations or stromal cells correlates with decreased apoptosis and increased mitogenic cell-state in colonic organoids (Fig. 5d).

**Fig. 6.**
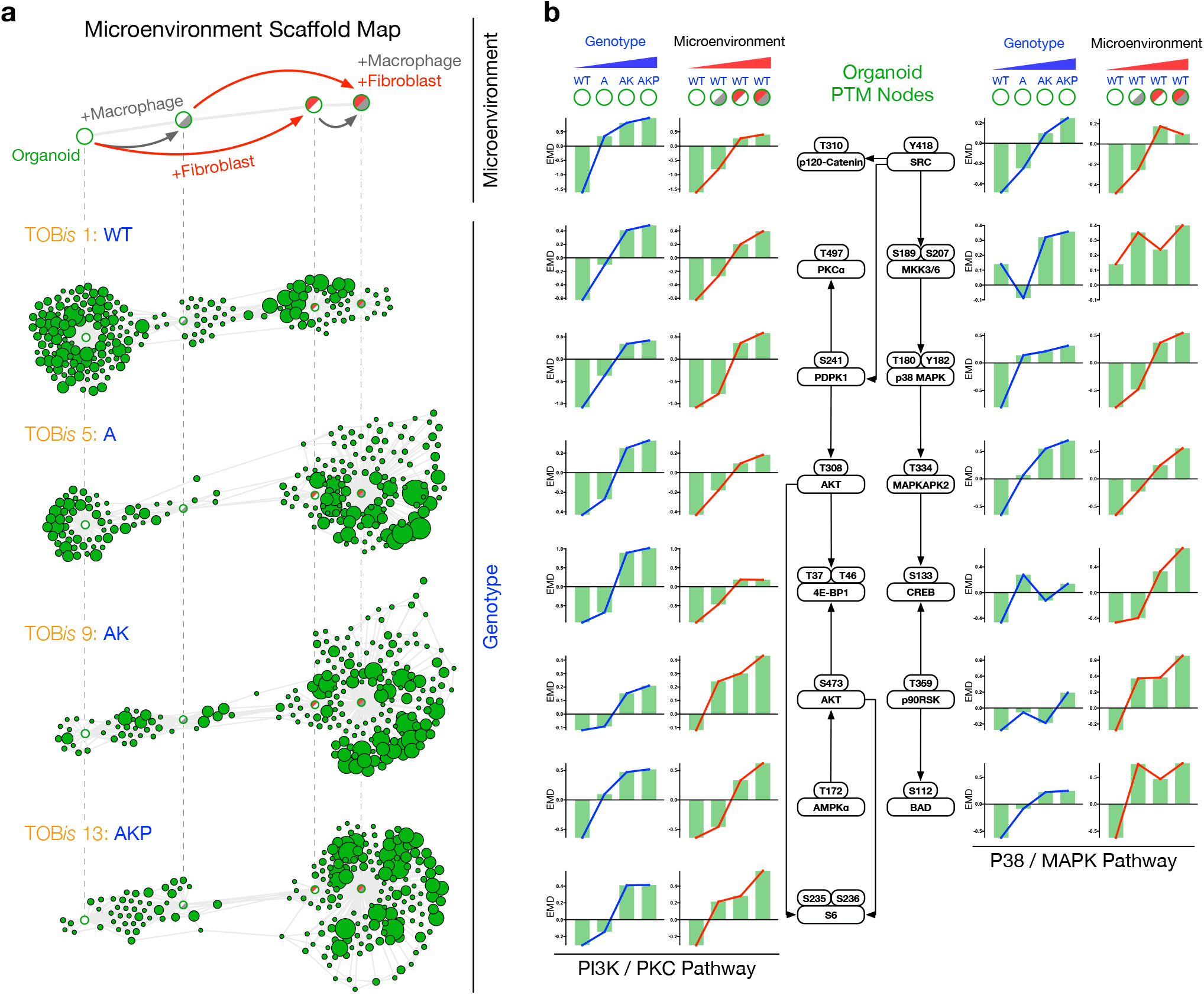
Oncogenic Mutations Mimic Stromal Signalling Networks. **a)** Scaffold maps constructed from WT organoids either alone or co-cultured with colonic fibroblasts and/or macrophages. Unsupervised distribution of A, AK, and AKP colonic organoids revealed that oncogenic mutations mimic signalling profiles driven by stromal fibroblasts and macrophages. (See Supplementary Fig. 9a for all genotype / microenvironment combinations and Supplementary Fig. 9b for mutation-driven Scaffold maps.) **b)** PTM-EMDs for PI3K / PKC and P38 / MAPK signalling nodes in colonic organoids following genotypic and microenvironmental regulation. Cell-type specific PTM analysis demonstrates oncogenic mutations and microenvironmental cues upregulate analogous signalling nodes in epithelial colonic organoids.

Taken together, multivariate cell-type specific PTM analysis of organoid co-cultures elucidated several fundamental processes in CRC: 1) oncogenic mutations re-structure signalling networks in cancer cells, whereas microenvironmental cues drive acute signalling flux, 2) stromal cells hyperactivate PI3K signalling in colonic epithelial cells that already carry *Kras* and *Trp53* mutations, and 3) oncogenic mutations cell-autonomously mimic an epithelial signalling state normally induced by stromal cells. These results collectively confirmed that TOB*is*-multiplexed MC enables discoveries of novel cell-type specific signalling relationships between different cell-types in organoid models of the tumour microenvironment.

## DISCUSSION

Organoids are heterocellular systems that comprise multiple cell-types and cell-states. Cell-type specific PTM signalling networks regulate major biological processes and are frequently dysregulated in disease. As a result, understanding cell-type specific signalling networks is fundamental to the utility of organoids and organoid co-cultures. Existing bulk PTM technologies (e.g. LC-MS/MS and anti-phospho antibody arrays) cannot describe cell-type or cell-state specific signalling relationships and therefore limit our understanding of organoid biology (9). While scRNA-seq can characterise cell-type specific transcription, it cannot measure protein-level signal transduction which ultimately drives biological phenotypes. To overcome these challenges, we demonstrated how a modified MC workflow that combines monoisotopic cell-type, cell-state, and PTM probes can be used to study cell-type specific signalling networks in organoids. This method uncovered novel cell-type specific signalling in intestinal epithelia and revealed an intimate relationship between cell-signalling and cell-state in organoids. We showed how <u>T</u>hiol-reactive <u>O</u>rganoid <u>B</u>arcoding *<u>i</u>n <u>s</u>itu* (TOB*is*) enables high-throughput comparison of signalling networks across different organoid mono- and co-cultures. Application of this technology to CRC TME organoid co-cultures revealed that oncogenic mutations mimic stromal signalling cues and demonstrated how highly mutated CRC cells can be further dysregulated by fibroblasts and macrophages.

While this study has focused on intestinal organoids, we expect this method to be fully compatible with organoids derived from other tissues (e.g. brain, liver, pancreas, kidney etc.). Cell-type identification probes for each tissue should be carefully validated, but otherwise the TOB*is* multiplexing and PTM analysis framework we report should be compatible with all organoid models (including those grown in defined hydrogels (37)). Moreover, our extension of MC to study colonic fibroblasts and macrophages implies that PTM signalling can be measured in any cell-type co-cultured with organoids (e.g. PBMCs co-cultured with organoids (3) and air-liquid interface tumour microenvironment organoids (38)).

In addition to standard single-cell organoid signalling experiments, the new barcoding technology reported here holds substantial promise for organoid screening. While drug screens of patient-derived organoid (PDO) monocultures have shown great potential (39, 40), their reliance on bulk viability measurements (e.g. CellTiter-Blue) implies that they cannot be used to evaluate drugs targeting stromal and/or immune cells or provide any mechanistic understanding of drug performance and/or resistance. In contrast, a TOB*is*-multiplexed MC drug / CRISPR screen will characterise cell-type specific signalling networks, cell-cycle states, and apoptotic readouts at single-cell resolution across all cell-types in PDO and PDO co-cultures. Given its ability to resolve multiple cell-types, TOB*is* MC would be particularly powerful for evaluating biological therapies against solid tumours where cell-type specificity is essential for resolving drug (e.g. CAR T-cell) and target (organoid) phenotypes. Future development of TOB*is* barcodes using additional TeMal (×7 possible) and Cisplatin (×4 possible) isotopologs will greatly expand organoid multiplexing capacity and advance this technology to high-throughput organoid screening applications.

In summary, this study demonstrates how a modified MC platform can reveal cell-type specific signalling networks in organoid monocultures and uncover novel cell-cell signalling relationships in organoid co-cultures. Given the widespread adoption of organoids as biomimetic models of healthy and diseased tissue, we propose cell-type specific PTM analysis as a powerful technology for multivariate organoid phenotyping.

## METHODS

Methods, including statements of data availability and any associated accession codes and references, are available as supplementary information on bioRxiv.

## Supporting information

Supplementary Table 5. Parameters for Single-Cell Signalling Data Analysis

## ACKNOWLEDGEMENTS

We are extremely grateful to Dr. Lukas Dow (Cornell University) for sharing CRC organoids, Dr. Xin Lu (University of Oxford) for sharing mouse intestines, and Dr. Olga Ornatsky (Fluidigm) for providing monoisotopic Cisplatin (^195^Pt and ^196^Pt). We thank the UCL CI Flow-Core for MC support and Dr. Leland McInnes (Tutte Institute) for UMAP advise. Graphical organoid renders were designed by Dr. Jeroen Claus (Phospho Biomedical Animation). This work was supported by Cancer Research UK (C60693/A23783), UCLH Biomedical Research Centre (BRC422), The Royal Society (RSG\Rd1\180234), and Rosetrees Trust (A1989).

## AUTHOR CONTRIBUTIONS

X.Q. Designed the study, performed organoid and MC experiments, analysed the data, and wrote the paper.

J.S. Developed TOB*is*, designed rare-earth metal antibody panels, performed MC analysis, and analysed data.

P.V. Isolated and characterised colonic fibroblasts, macrophages, cultured organoids, and analysed data.

P.K. Performed UMAP, EMD, DREMI, and PCA data analysis.

M.N. Developed TeMal barcodes.

S.A. Provided murine monocytes and intestines for fibroblast isolation.

V.L. Provided murine small intestines for organoid isolation.

C.T. Designed the study, analysed the data, and wrote the paper.

## COMPETING INTERESTS

M.N. has pending intellectual property on the use of tellurium reagents for mass cytometry applications which has been licensed to Fluidigm Corporation.

