## Supplementary Methods and Figures for "Single-Cell Signalling Analysis of Heterocellular Organoids"

#### Organoid Culture

Intestinal organoids were generated as describe by Sato *et al.*[1]. Briefly, the small intestine of 8- to 12-week-old *Lgr5-EGFP-ires-CreERT2* mice was dissected, opened longitudinally, and cut into 2- to 5-mm segments. Tissue fragments were washed with ice-cold PBS and incubated with 2 mM EDTA (Sigma 03690) in PBS (Thermo 10010056) for 1 hr at 4 °C. After removal of EDTA, tissue fragments were washed vigorously in cold PBS to release the crypts. Supernatant fractions from the washes were collected and centrifuged at 1,200 rpm for 5 mins. Cells were washed with 15 mL advanced DMEM/F-12 (Thermo 12634010), passed through a 70 µm cell strainer (Fisher 11597522) to enrich for intestinal crypts, and centrifuged at 600 rpm for 2 mins. The cell pellet was resuspended in Growth Factor Reduced Matrigel (Corning 354230) and cultured at 37 °C in the presence of 5% $CO_2$.

Small intestinal organoids were maintained in advanced DMEM/F-12 (Thermo 12634010) supplemented with 2 mM L-Glutamine (Thermo 25030081), 1 mM N-Acetyl-L-Cysteine (Sigma A9165), 10 mM HEPES (Sigma H3375), 1× B-27 Supplement (Thermo 17504044), 1× N-2 Supplement (Thermo 17502048), 50 ng/mL murine EGF (mEGF, Thermo PMG8041), 50 ng/mL murine Noggin (mNoggin, Peprotech 250-38), 500 ng/mL murine R-Spondin-1 (mR-spondin-1, Peprotech 315-32), and 1× HyClone™ Penicillin Streptomycin Solution (Fisher SV30010).

Murine colorectal cancer (CRC) organoids carrying oncogenic mutations (*shApc* (A), *shApc* and $Kras^{G12D/+}$ (AK), or *shApc*, $Kras^{G12D/+}$, and $Trp53^{R172H/-}$ (AKP)[2, 3]) were a kind gift from Prof. Lukas Dow (Cornell University) and are described in Dow & O'Rourke et al., Cell, 2015[2] and O'Rourke et al., Nature Biotechnology, 2017[3]. Colonic organoids were maintained in advanced DMEM/F-12 (Thermo 12634010) supplemented with 2 mM L-Glutamine (Thermo 25030081), 1 mM N-Acetyl-L-Cysteine (Sigma A9165), 10 mM HEPES (Sigma H3375), 1× B-27 Supplement (Thermo 17504044), 1× N-2 Supplement (Thermo 17502048), 100 ng/mL murine WNT-3a (mWNT-3a, Peprotech 315-20), 50 ng/mL murine EGF (mEGF, Thermo PMG8041), 50 ng/mL murine Noggin (mNoggin, Peprotech 250-38), 500 ng/mL murine R-Spondin-1 (mR-spondin-1, Peprotech 315-32), 10 mM Nicotinamide (Sigma N0636), and 1× HyClone™ Penicillin Streptomycin Solution (Fisher SV30010). For passaging, small intestinal organoids were retrieved from Matrigel using ice-cold PBS and broken up mechanically by passing through a 23G, 5/8'' needle (Terumo AN-2316R). Colonic organoids were dissociated mechanically by pipetting or enzymatically with TrypLE™ Express Enzyme (Thermo 12604013). Organoid fragments were collected using a benchtop centrifuge, washed with ice-cold PBS, and reseeded in fresh Matrigel. The passage was performed every 4 to 7 days at a ratio of 1:2 or 1:3.

**Pre-Treatment and Fixation of Organoids for Mass Cytometry**

$^{127}$5-Iodo-2′-deoxyuridine ($^{127}$IdU) (Fluidigm 201127) was added directly to organoid culture media to a final concentration of 25 μM and incubated at 37 °C for 30 mins before fixation to identify cells in S-phase[4]. 5 mins before fixation, protease and phosphatase inhibitors (Sigma P8340 / Sigma 4906845001) were added to organoid cultures to preserve cell signalling during fixation[5]. As dissociation of live tissue alters cellular states[6] (including PTMs[5]), all organoids were fixed in 4% PFA (Thermo J19943K2) for 60 mins at 37 °C to preserve cell-signalling. (Note: during method optimisation, we also trialled alternative fixatives such as Glutaraldehyde (0.2%, 1%, 2.5%), Ethanol (5%, 10%), and Glyoxal (pH 4.0, pH 5.0), but concluded that 4% PFA was optimal. 1.6% PFA can also be used but we encourage users to test a range of PFA for their specific antibody panel.) Following fixation, organoids were washed ×2 with PBS and incubated in 250 nM $^{194/8}$Cisplatin (Fluidigm 201194 / 8) in PBS for 10 mins on a rocker to stain dead cells[7]. During optimisation we found this condition yields strong $^{194/8}$Pt staining with a wide dynamic range suitable for efficient dead cell removal *in silico*. Organoids were then washed ×2 with PBS to remove residual Cisplatin. Fixed organoids were subsequently dissociated for single-cell analysis immediately or stored at 4 °C.

**Single-Cell Dissociation of Fixed Organoids**

After organoids were fixed and stained with Cisplatin, the final wash was removed and a solution of fresh 0.5 mg/mL Dispase II (Thermo 17105041), 0.2 mg/mL Collagenase IV (Thermo 17104019), and 0.2 mg/mL DNase I (Sigma DN25) in PBS was added to the organoids. (During optimisation we found Dispase II is essential for disrupting epithelial cell-cell contacts, Collagenase IV improves Matrigel degradation, and DNase I digests extracellular genomic DNA from dead organoid cells to reduce sample viscosity. We encourage users to test alternative dissociation enzymes for the specific cellular composition of their experimental system.) Organoid droplets were then scraped from the well, pooled, and the enzyme / organoid solution was transferred to a gentleMACS C-Tube (Miltenyi 130-096-334). Fixed organoids were dissociated into a single-cell suspension using the gentleMACS Octo Dissociator (with Heaters) (Miltenyi 130-096-427) at 37 °C for 50 mins using a custom program. Following dissociation, C-Tubes were centrifuged at 800 ×g for 1 min to collect cells from blades and all liquid was transferred to a fresh polypropylene FACS tube (Corning 352063). Single organoid cells were then washed ×2 in Cell Staining Buffer

(CSB) (Fluidigm 201068) (5 mins, 800 ×g) to remove enzymes / cellular debris and 35 µm filtered (Fisher 10585801) (70 µm (Fisher 11597522) when the culture contains fibroblasts) to remove residual clumps.

**Heavy-Metal Antibody Conjugation and Panel Design**

All antibodies were custom conjugated[8] with rare-earth / heavy metals using X8 polymers and monoisotopic metals from Fluidigm (Fluidigm 201300). Non-Fluidigm metals / nitrates were also used: $^{89}$Y (Sigma 217239), $^{113}$In (Trace Sciences), $^{115}$In (Trace Sciences), $^{157}$Gd (Trace Sciences), and $^{209}$Bi (Sigma 254150)[9]. Antibody panels (Supplementary Tables 1 and 2) were carefully designed and titrated in accordance with known monoisotopic impurities[10] and antigen abundance to ensure minimal cross-channel contamination.

**Mass Cytometry Analysis of Single Organoid Cells**

1–5 × $10^6$ fixed single organoid cells were blocked in CSB and stained with organoid-specific extracellular rare-earth metal antibody cocktails (Supplementary Tables 1 and 2) for 30 mins. Cells were then washed ×2 with CSB (5 mins, 800 ×g) and permeabilised in 0.1% Triton X-100 (Sigma T8787) in PBS for 30 mins. Cells were washed ×2 in CSB and further permeabilised with ice-cold 50% methanol (Fisher 10675112) for 10 mins on ice. (Note: during method optimisation we found dual 0.1% Triton X-100 and 50% Methanol provides the best all-round permeabilisation for a broad range of anti-PTM antibodies.) Permeabilised cells were then washed ×2 in CSB and stained with intracellular rare-earth metal antibody cocktails (Supplementary Tables 1 and 2) for 30 mins. Stained cells were washed ×2 in CSB, fixed in fresh 1.6% formaldehyde (Thermo 28906) for 10 mins, washed in CSB, and incubated in DNA Intercalator (Fluidigm 201192A) overnight at 4 °C. The following day, cells were washed ×2 in CSB, resuspended in Maxpar Water (Fluidigm 201069) containing 20% (v/v) EQ Beads[11] (Fluidigm 201078) and 2 mM EDTA at ~0.5 × $10^6$ cells / mL. Cells were then 35 μm filtered (Fisher 10585801) (70 μm (Fisher 11597522) when the culture contains fibroblasts) and immediately analysed using a Helios Mass Cytometer (Fluidigm) (100 – 300 events / sec). Files were normalised against EQ beads, de-barcoded[12] into each experimental condition (when required), and uploaded to the Cytobank platform (http://www.cytobank.org/).

**Immunofluorescence (IF) Staining of Organoids**

Intestinal organoids were cultured in 8-well *μ*-Slides (ibidi 80826). After culture medium was removed, cells were washed with PBS and fixed with 4% PFA (Thermo J19943K2) for 30 mins at 4 °C. Cells were washed twice with PBS and permeabilised with 0.2% Triton™ X-100 (Sigma T8787) in PBS for 30 mins at room temperature. Cells were washed again, incubated with PBS containing 1% BSA (CST #9998) and 0.3% Triton™ X-100 for 30 mins, followed by incubation with primary antibodies diluted in 1% BSA / 0.3% Triton™ X-100 / PBS overnight at 4 °C. Cells were washed with PBS, stained with secondary antibodies and 4',6-Diamidino-2-Phenylindole (DAPI) (Thermo D1306), with or without Alexa Fluor™ 488 / 568 Phalloidin (Thermo A12379 / A12380) for 1 hr at room temperature, away from light. Cells were washed with PBS and mounted with Fluoromount-G™ mounting medium (Thermo 00-4958-02). Samples were imaged with a Zeiss LSM880 confocal microscope and images were analysed using FIJI[13]. EdU staining was performed using the Click-iT Plus EdU Alexa Fluor 647 Imaging Kit (Thermo C10640) following the manufacture's protocol.

**Small Intestinal Organoid Directed Differentiation**

Small intestinal organoids were seeded and cultured in complete organoid medium (described above) for 24 hrs to allow organoid recovery, and treated with combinations of 3 µM CHIR99021 (GSK-3β inhibitor) (Cambridge Bioscience SM13), 2 µM IWP-2 (PORCN inhibitor) (Cambridge Bioscience 13033), 1 mM Valproic Acid (HDAC inhibitor) (Cambridge Bioscience SM39), and 10 µM DAPT (γ-Secretase inhibitor) (Cambridge Bioscience SM15) for 3 days to direct organoid differentiation towards specific cell-types as described in Yin et al.[14] (Supplementary Fig. 1). Directed-differentiated organoids were analysed by IF and MC as described above.

**Single-Cell Dissociation of Murine Intestinal Crypts**

The small intestine of 8- to 12-week-old *Lgr5-EGFP-ires-CreERT2* mice was dissected and intestinal crypts were isolated as described above (see 'Organoid Culture'). The crypts were resuspended in 5 mL of TrypLE™ Express Enzyme and incubated at 37 °C for 45 mins, mixed every 10 mins to avoid cells clumping. The cells were centrifuged at 1,200 rpm for 5 mins, resuspended in 5 mL of 4% PFA, and fixed at 37 °C for 1 hr. Fixed cells were washed once with PBS, 35 µm filtered twice to remove residual clumps, and stored at 4 °C prior to MC analysis.

**Amine- versus Thiol-Reactive *in situ* Organoid Probes**

To investigate alternative probe chemistries for *in situ* organoid barcoding, fixed small intestinal organoids were stained with either 50 nM Alexa Fluor™ 647 NHS ester (Thermo A20006) or 50 nM Alexa Fluor™ 647 $C_2$ maleimide (Thermo A20347) for 1 hr while still in Matrigel. Organoid probe intensities were visualised by confocal microscopy using identical settings for each probe (as described above). To confirm

IF observations using MC, fixed small intestinal organoids were stained with either 200 nM NHS-DOTA (Macrocyclics B-280) or 200 nM maleimide-DOTA (Macrocyclics B-272) coupled to $^{157}$Gd (Trace Sciences) for 1 hr *in situ* (in Matrigel) or *ex situ* (removed from Matrigel). Organoids were then dissociated into single cells and analysed by MC (as described above).

**Thiol-reactive Organoid Barcoding *in situ* (TOB*is*)**

Fixed organoids were washed ×2 in PBS and stained *in situ* with 10 nM Cisplatin ($^{196}$Pt, $^{198}$Pt)[15] (a kind gift from Dr. Olga Ornatsky, Fluidigm) and 1—3 µM TeMal ($^{124}$Te, $^{126}$Te, $^{128}$Te, $^{130}$Te)[16] barcodes for 1 hr on a rocker (barcoding matrices in Supplementary Tables 3 and 4). (Note: organoids can also be barcoded overnight at 4 °C.) The lower concentration of Cisplatin used for TOB*is* (10 nM) relative to dead cell stains (250 nM) is to ensure $^{196}$Pt and $^{198}$Pt signal intensities align with $^{124}$Te, $^{126}$Te, $^{128}$Te, and $^{130}$Te signals. TOB*is* barcoding can be performed in 6-well (3 mL barcodes), 12-well (2 mL barcodes), 24-well (1 mL barcodes), 48-well (500 µL barcodes), and 96-well (200 µL barcodes) plates. In our experience up to 10,000 cells / µL Matrigel (representing confluent intestinal organoid cultures) can be efficiently barcoded, but we suggest optimising barcode concentrations for alternative model systems. Organoids were washed ×3 in CSB containing 1 mM L-Glutathione (Sigma G6529) for 5 mins on a rocker to quench unbound barcodes (Supplementary Fig. 5h) and ×1 in PBS prior to pooled-dissociation (described above).

To directly compare TOB*is* and Maxpar (Fluidigm 201060) performance for *in situ* barcoding, 20 wells of established small intestinal organoids in the 12-well culture format (3 × 30 µL Matrigel droplets / well) were fixed as described above and washed ×2 with PBS. TOB*is* Barcode 1—20 was mixed respectively with Maxpar Barcode 1—

20 in PBS and added to each well. The cells were incubated at room temperature for 60 mins, washed ×3 in CSB containing 1 mM L-Glutathione (Sigma G6529) for 5 mins on a rocker to quench unbound barcodes, dissociated into single cells, and analysed by MC.

**Maxpar *in situ* versus ex si*tu* Organoid Barcoding Comparison**

Established small intestinal organoids were fixed in 4% PFA and washed ×2 with PBS. Maxpar barcodes were resuspended in Barcode Perm Buffer as per the manufacturer's protocol (Fluidigm 201060). For *in situ* barcoding, cells were incubated in Barcode Perm Buffer for 10 mins followed by barcoding solution incubation at room temperature for 60 mins. The cells were washed ×2 with CSB to quench unbound barcodes and proceeded to dissociation as described above. For *ex situ* barcoding, cells were dissociated and barcoded according to the manufacturer's protocol. The *in situ* and *ex situ* samples were pooled into a single tube and analysed by MC. Cells were de-barcoded and single-cell counts were analysed in GraphPad Prism 7 (two-tailed unpaired *t*-test).

To demonstrate the reactivity of Maxpar barcodes to Matrigel (hence its incompatibility with organoid barcoding *in situ*), empty Matrigel droplets were seeded in 12-well plates (3 × 30 µL droplets / well), fixed with 4% PFA, washed ×2 with PBS, and incubated with PBS, Maxpar barcode #20 ($^{106}$Pd, $^{108}$Pd, and $^{110}$Pd) resuspended in PBS or CSB for 60 mins at room temperature. The Matrigel droplets were then washed ×3 with CSB, ×3 with PBS, and dissolved in ice-cold Maxpar Water (Fluidigm 201069). Matrigel concentration was measured by BCA assay (Thermo 23225) and all samples were diluted to a protein concentration of 250 µg/mL prior to analysis on solution mode using a Helios Mass Cytometer (time per reading = 1 sec, settling time

= 10 msec). The dual counts of $^{110}$Pd were measured and analysed in GraphPad Prism 7 (two-tailed unpaired *t*-test).

**TOB*is* versus Maxpar Cell-Recovery Comparison**

During optimisation we observed that organoids barcoded using the Maxpar Cell-ID™ 20-Plex Pd Barcoding Kit (Fluidigm 201060) had much smaller cell pellets than those processed by TOB*is*. We hypothesised this cell-loss was due to the increased dissociation and centrifugation steps required for Maxpar barcoding when compared to TOB*is*. To investigate this, we directly compared both TOB*is* and Maxpar barcoding recoveries across a range of organoid seeding densities.

Established small intestinal organoids (4-day-culture) were removed from Matrigel and reseeded in 24-well plates (1 × 50 µL Matrigel droplet / well) across 6 serial dilutions (100—3.125%) in duplicate to generate a dynamic range of organoid cell numbers. After recovering for 12 hrs in complete organoid culture medium, organoids were fixed in 4% PFA and washed in PBS (see above). One replicate of the organoids was barcoded *in situ* using TOB*is* (described above) and the other replicate was individually dissociated and barcoded *ex situ* using the Maxpar kit following the manufacturer's instructions. Cells were stained with rare-earth metal antibodies and analysed by MC. Cells were de-barcoded and single-cell counts were analysed in GraphPad Prism 7 (two-tailed ratio-paired *t*-test).

To investigate organoid cell-type recovery between Maxpar and TOB*is* barcoding, equal cell numbers across replicates of each strategy were identically gated in UMAP space (see below) and the percentages of stem, Paneth, enteroendocrine, tuft, goblet cells, and enterocytes were calculated. Cell-type percentages were analysed in GraphPad Prism 7 (linear regression of correlation).

**Small Intestinal Organoid Time-Course**

Intestinal organoids were retrieved from Matrigel using ice-cold PBS, broken up mechanically by sequentially passing through a 23G, 5/8” needle (Terumo AN-2316R) 6 times and a 26G, 1/2” needle (HSW, 4710004512) 3 times to obtain a uniform cell suspension. Organoid fragments were 70 µm filtered twice and centrifuged at 200 ×g for 5 mins. The cell pellet enriched with single crypts was washed with cold PBS, collected using a benchtop centrifuge, and resuspended in Matrigel prior to seeding. The cell ratio seeded for Days 1—7 was 30: 9: 6: 5: 4: 3: 3 to ensure comparable organoid recovery and density from each time point. At each time point, the organoids were incubated with 25 µM $^{127}$IdU, protease / phosphatase inhibitors, and fixed in 4% PFA for 60 mins at 37 °C as described above. Organoids were washed with PBS and stored at 4 °C until samples from all time points were collected. All samples were stained with 250 nM $^{194}$Cisplatin, TOB*is* barcoded (described above) (Supplementary Table 3), stained, and analysed in one MC experiment (Supplementary Table 1, 50 parameters (40 antibodies) / cell).

**Colonic Fibroblast Isolation, Immortalisation, and Cell Culture**

Colonic fibroblasts were isolated as described by Khalil et al.[17]. Freshly dissected murine (C57BL/6, 6- to 8-week-old) colon tissue was flushed with ice-cold PBS, cut open, washed again in PBS, and incubated in 5 mM EDTA / PBS at 250 rpm, 37 °C for 15 mins. This process was repeated for a total of ×5 EDTA / PBS washes. Washed colon tissue was transferred to a fresh tube of sterile DMEM (Thermo 41966052) supplemented with 1 mg/mL Dispase II (Thermo 17105041) and 1 mg/mL Collagenase D (Sigma 11088858001). The colon / enzyme solution was incubated at 250 rpm, 37 °C for 30—60 mins (until the tissue started to look ‘stringy’). Digested colon tissue was then centrifuged at 200 ×g, 4 °C for 5 mins. Supernatant was discarded and the pellet was resuspended in 10 mL ACK Lysing Buffer (Thermo A1049201). Cells were centrifuged at 200 ×g, 4 °C for 5 mins, and the pellet was resuspended in DMEM + 10% FBS (Thermo 10082147). Cells were 100 µm filtered (Miltenyi 130-098-463) into a T75 flask and incubated at 5% $CO_2$, 37 °C. After 3 hrs, cells were washed ×2 with PBS to remove debris. Adhered cells were cultured with DMEM + 10% FBS + 1× Insulin-Transferrin-Selenium (ITS-G) (Thermo 41400045). After 1 week of culture, fibroblasts were observed to proliferate, while other cell-types (e.g. epithelial cells and leukocytes) senesce and/or die. Colonic fibroblasts were immortalised using pBABE-HPV-E6 retrovirus produced in Phoenix-ECO cells (a kind gift from Prof. Erik Sahai, The Francis Crick Institute, London) and stably transfected with RFP using the pCMV-DsRed-Express plasmid with Lipofectamine 3000 (Thermo L3000001) to aid co-culture visualisation. Immortalised colonic fibroblasts were cultured in DMEM + 10% FBS + 1× ITS-G at 5% $CO_2$, 37 °C. Cells were checked for mycoplasma infection monthly using the MycoAlert™ PLUS Mycoplasma Detection Kit (Lonza LT07-701) and remained negative throughout this project. IF staining confirmed that colonic fibroblasts were positive for intestinal mesenchymal markers such as Vimentin (D21H3, CST), Podoplanin (PDPN) (8.1.1, BioLegend), PDGFRα (APA5, Abcam), FOXL1 (ab95286, Abcam), and GLI-1 (C-1, Santa Cruz) in both 2D and 3D cultures.

**Primary Macrophage Isolation and Cell Culture**

Freshly dissected murine (C57BL/6, 10- to 12-week-old females) femurs and tibias (Charles River Laboratories) were flushed ×5 with 10 mL RPMI 1640 Medium (Thermo 11875093) + 10% FBS (Thermo 10082147). Cells were centrifuged at 300 ×g for 5 mins, resuspended in RPMI + 10% FBS, 40 µm filtered (Fisher 11587522), and centrifuged at 300 ×g for 5 mins. Supernatant was discarded and the pellet was resuspended in 2 mL ACK Lysing Buffer (Thermo A1049201) for 5 mins at room temperature. Monocytes were washed in PBS, centrifuged at 300 ×g for 5 mins, resuspended in 1 mL Recovery™ Cell Culture Freezing Medium (Thermo 12648010), and stored in liquid nitrogen until use. Bone marrow-derived macrophages were expanded and activated in RPMI + 10% FBS + 25% L929-cell conditioned media (LCCM) before experiments. IF staining confirmed that the cells were positive for intestinal macrophages markers such as CD45 (30-F11, BioLegend), CD68 (FA-11, BioLegend), CD11b (M1/70, BioLegend), F4/80 (BM8, BioLegend), and CX3CR1 (SA011F11, BioLegend) in both 2D and 3D cultures.

**Heterocellular CRC Tumour Microenvironment (TME) Organoid Culture**

Wild-type (WT) murine colonic organoids and CRC organoids carrying oncogenic mutations A, AK, and AKP were a kind gift from Prof. Lukas Dow (Cornell University) and cultured as described above. Following expansion in complete media, organoids were cultured in the absence of exogenous growth factors (mEGF, mNoggin, mR-Spondin-1 and mWnt-3a — WENR) for 8 hrs prior to the experiment. Colonic fibroblasts were cultured in DMEM supplemented with reduced FBS (2%) and 1× ITS-G for 24 hrs before the experiment. Primary bone marrows were differentiated into macrophages using RPMI + 10% FBS + 25% LCCM for 7 days before the experiment. To establish the CRC TME culture, organoids were passaged at a ratio of ~1:2.5; colonic fibroblasts were seeded at 6,000 cells / µL, 5,000 cells / µL and 4,000 cells / µL Matrigel for monoculture, 2-way co-cultures, and 3-way co-cultures respectively; primary macrophages were seeded at 9,000 cells / µL, 8,000 cells / µL and 7,000 cells / µL Matrigel for monoculture, 2-way co-cultures, and 3-way co-cultures respectively. Organoids, fibroblasts, and macrophages were mixed in Matrigel before seeding at 3 × 30 µL droplets per well in a 12-well plate. Each microenvironment culture was maintained in WENR-free advanced DMEM/F-12 (Thermo 12634010) supplemented with 2 mM L-Glutamine (Thermo 25030081), 1 mM N-Acetyl-L-Cysteine (Sigma A9165), 10 mM HEPES (Sigma H3375), 1× B-27 Supplement (Thermo 17504044), 1× N-2 Supplement (Thermo 17502048), 1× Insulin-Transferrin-Selenium-Sodium Pyruvate (ITS-A) (Thermo 51300044), and 1× HyClone™ Penicillin Streptomycin Solution (Fisher SV30010) for 48 hrs. All cultures were incubated with 25 µM $^{127}$IdU, protease / phosphatase inhibitors, and fixed in 4% PFA for 60 mins at 37 °C (as described above). Dead cells were stained with 250 nM $^{194}$Cisplatin as described above. Organoids were barcoded using TOB*is* (Supplementary Table 4), pooled into a single tube, dissociated into single cells, 70 µm filtered, and stained for MC analysis (Supplementary Table 2, 50 parameters (40 antibodies) / cell). Single organoid, fibroblast, and macrophage cells were analysed by MC (as described above).

**Single-Cell Signalling Data Analysis**

All single cells were gated for Gaussian parameters (Event length, Centre, Residual, and Width values), DNA$^{high}$ ($^{191}$Ir and $^{193}$Ir), and Cisplatin$^{low}$ ($^{194/8}$Pt). For small intestinal organoids and colonic organoids, intact epithelial cells were gated with EpCAM$^{+}$ / Pan-CK$^{+}$ and CEACAM1$^{+}$ / Pan-CK$^{+}$ respectively. Intact colonic fibroblasts were gated with RFP$^{+}$ / PDPN$^{+}$, and primary macrophages were gated with CD68$^{+}$ / F4/80$^{+}$. Removal of cells stained positive for mutually exclusive cell-type / cell-state markers was performed as part of data pre-processing procedures (gating strategies incorporated in the publicly deposited datasets). Cells were then clustered and visualised in UMAP (Uniform Manifold Approximation and Projection)[18] space and gated for cell-type and cell-state makers before proceeding to PTM analysis (Supplementary Fig. 2a—c).

UMAP analysis was performed with the Python package *umap* (https://umap-learn.readthedocs.io/en/latest) using default parameters unless otherwise specified (Supplementary Table 5). UMAP was used to visualise high-dimensional MC datasets in two-dimensional space, where cell gating was performed to identify cell populations or to remove residual outliers when required. All data was arcsinh transformed with a cofactor of 5. All parameters used to generate UMAPs are listed in Supplementary Table 5.

Earth Mover's Distance (EMD) was computed with the Python package *scprep*[19] (https://github.com/KrishnaswamyLab/scprep) using default parameters. EMD scores were signed by the difference of the median intensity of a given parameter between the population of interest relative to the denominator (specified below): positive for up-regulation or negative for down-regulation. Cell populations were manually gated and exported from Cytobank, with all channels to be analysed arcsinh transformed (cofactor = 5). For single-time-point small intestinal organoids (Fig. 2b and Supplementary Fig. 3), EMD was calculated between each cell-type / state and the entire epithelial cell population. For the small intestinal organoid time-course experiment (Fig. 4c), EMD was calculated between each cell-type from each time point against the combined population of all epithelial cells across all time points. For the CRC TME model (Figs. 5d, e, 6b and Supplementary Figs. 7, 8a—d), EMD was calculated between each cell-type in each condition and the combined population of all cell-types across all conditions.

*k*-Nearest Neighbours Density Resampled Estimation of Mutual Information (*k*NN-DREMI)[20] was computed with the Python package *scprep*[19] using default parameters. Cell populations were manually gated in UMAP space and exported from Cytobank, with all the channels to be analysed arcsinh transformed (cofactor = 5). For small intestinal organoids (Fig. 2b and Supplementary Fig. 3), *k*NN-DREMI scores among 28 PTMs (Supplementary Table 1) were calculated, which yielded a total of 756 PTM-PTM combinations for each cell-type / cell-state. Similarly, *k*NN-DREMI scores for 756 PTM-PTM pairs across 28 PTMs (Supplementary Table 2) were computed for each cell-type in the CRC TME model (Fig. 5f, Supplementary Figs. 7, 8a, b, e, and f). Heatmaps were generated using the R package *RColorBrewer* (https://cran.r-project.org/web/packages/RColorBrewer/) based on EMD and DREMI calculations. Signalling maps were compiled in OmniGraffle Professional from the heatmaps with the nodes (PTMs) coloured by EMD scores and edges (PTM-PTM pairs) by DREMI scores.

Principal Component Analysis (PCA) was performed on z-score normalised EMD (Fig. 2c, 5e, and Supplementary Fig. 8c, d) or DREMI (Fig. 5f and Supplementary Fig. 8e, f) scores using the Scikit-Learn package *PCA estimator* in Python (https://scikit-learn.org/stable/modules/generated/sklearn.decomposition.PCA.html) with default parameters. All measurements used for PCA are listed in Supplementary Table 5.

Force-directed Scaffold Maps[21] were constructed using the R package *Scaffold* (https://github.com/nolanlab/scaffold) with parameters specified below. Landmark populations were manually gated and exported from Cytobank with all data arcsinh transformed (cofactor = 5). For small intestinal organoid directed differentiation (Supplementary Fig. 1), pRB [S807/S811]$^{+}$ cells were used for the Scaffold analysis. UMAP-gated stem, Paneth, enteroendocrine, goblet, tuft cells, and enterocytes from untreated organoids were used as landmark nodes, and the untreated sample was used as the reference dataset. All measurements including cell-type / cell-state markers and PTMs were used to generate the Scaffold maps (Supplementary Table 5). For the CRC TME model (Fig. 6a and Supplementary Fig. 9), epithelial cells from each condition were clustered by the measurements of all cell-type / cell-state markers and PTMs (Supplementary Table 5). Microenvironment-focused Scaffold maps (Fig. 6a and Supplementary Fig. 9a) were generated using epithelial cells from WT organoid monoculture and WT organoids co-cultured with macrophages and/or fibroblasts as landmark nodes, and epithelial cells from the WT organoid / macrophage / fibroblast co-culture sample as the reference dataset. For the genotype-focused Scaffold maps (Supplementary Fig. 9b), epithelial cells from monocultures of WT, A, AK, AKP organoids were used as landmark nodes, and the WT sample was used as the reference dataset. Selected cell-type and cell-state markers were used to generate the Scaffold maps (Supplementary Table 5).

**Data Availability**

All raw data, processed data, and working illustrations are available as a Community Cytobank project (https://community.cytobank.org/cytobank/experiments#project-id=1271).

#### Supplementary Table 1 – Small Intestinal Organoid Mass Cytometry Panel (Figures 1 & 2)

| Isotope-Metal | Antigen / Target | Antibody Clone | Supplier |
|---|---|---|---|
| 89-Y | Phospho-Histone H3 [S28] | HTA28 | BioLegend |
| 113-In | CD326 (EpCAM) | G8.8 | BioLegend |
| 115-In | Pan-Cytokeratin (Pan-CK) | AE1/AE3 | BioLegend |
| 124-Te | TOB*is* Barcode | - | Prof. Mark Nitz |
| 126-Te | TOB*is* Barcode | - | Prof. Mark Nitz |
| 127-IdU | S-Phase | - | Fluidigm |
| 128-Te | TOB*is* Barcode | - | Prof. Mark Nitz |
| 130-Te | TOB*is* Barcode / TePhe | - | Prof. Mark Nitz |
| 141-Pr | Phospho-PDPK1 [S241] | J66-653.44.22 | BD Biosciences |
| 142-Nd | Cleaved-Caspase 3 [D175] | D3E9 | CST |
| 143-Nd | C-MYC | D84C12 | CST |
| 144-Nd | Lysozyme | BGN/06/961 | Abcam |
| 145-Nd | FABP1 | 328605 | R&D Systems |
| 146-Nd | Phospho-MKK4/SEK1 [S257] | C36C11 | CST |
| 147-Sm | Phospho-BTK [Y551] | 24a/BTK | BD Biosciences |
| 148-Nd | Phospho-SRC [Y418] | SC1T2M3 | BD Biosciences |
| 149-Sm | Phospho-4E-BP1 [T37/46] | 236B4 | CST |
| 150-Nd | Phospho-RB [S807/811] | J112-906 | BD Biosciences |
| 151-Eu | Phospho-PKCα [T497] | K14-984 | BD Biosciences |
| 152-Sm | Phospho-AKT [T308] | J1-223.371 | BD Biosciences |
| 153-Eu | Phospho-CREB [S133] | 87G3 | CST |
| 154-Sm | Phospho-SMAD1 [S463/465]<br>Phospho-SMAD5 [S463/465]<br>Phospho-SMAD9 [S465/467] | D5B10 | CST |
| 155-Gd | Phospho-AKT [S473] | D9E | CST |
| 156-Gd | Phospho-NF-κB p65 [S529] | K10-895.12.50 | BD Biosciences |
| 157-Gd | Phospho-MKK3 [S189] / MKK6 [S207] | D8E9 | CST |
| 158-Gd | Phospho-p38 MAPK [T180/Y182] | D3F9 | CST |
| 159-Tb | Phospho-MAPKAPK2 [T334] | 27B7 | CST |
| 160-Gd | Phospho-AMPKα [T172] | 40H9 | CST |
| 161-Dy | Phospho-BAD [S112] | 40A9 | CST |
| 162-Dy | LRIG1 | Polyclonal | R&D Systems |
| 163-Dy | Phospho-p90RSK [T359] | D1E9 | CST |
| 164-Dy | Phospho-p120-Catenin [T310] | 22/p120 (pT310) | BD Biosciences |
| 165-Ho | β-Catenin [Active] | D13A1 | CST |
| 166-Er | Phospho-GSK-3β [S9] | D85E12 | CST |
| 167-Er | Phospho-ERK1/2 [T202/Y204] | 20A | BD Biosciences |
| 168-Er | Phospho-SMAD2 [S465/467]<br>Phospho-SMAD3 [S423/425] | D27F4 | CST |
| 169-Tm | GFP | 5F12.4 | Fluidigm |
| 170-Er | Phospho-MEK1/2 [S221] | 166F8 | CST |
| 171-Yb | CLCA1 | EPR12254-88 | Abcam |
| 172-Yb | Phospho-S6 [S235/236] | D57.2.2E | CST |
| 173-Yb | DCAMKL1 | 6F9 | Sigma |
| 174-Yb | Chr-A | C-12 | Santa Cruz |
| 175-Lu | CD44 | IM7 | BioLegend |
| 176-Yb | Cyclin B1 | GNS-11 | BD Biosciences |
| 191-Ir | DNA | - | Fluidigm |
| 193-Ir | DNA | - | Fluidigm |
| 194-Pt | Dead Cells | - | Fluidigm |
| 196-Pt | TOB*is* Barcode | - | Fluidigm |
| 198-Pt | TOB*is* Barcode | - | Fluidigm |
| 209-Bi | Di-Methyl-Histone H3 [K4] | C64G9 | CST |

Extracellular
Intracellular
Non-Antibody Parameter

#### Supplementary Table 2 – CRC Tumour Microenvironment Mass Cytometry Panel (Figure 3)

| Isotope-Metal | Antigen / Target | Antibody Clone | Supplier |
|---|---|---|---|
| 89-Y | Phospho-Histone H3 [S28] | HTA28 | BioLegend |
| 113-In | CD66a (CEACAM1) | CC1 | Thermo |
| 115-In | Pan-Cytokeratin (Pan-CK) | AE1/AE3 | BioLegend |
| 124-Te | TOB*is* Barcode | - | Prof. Mark Nitz |
| 126-Te | TOB*is* Barcode | - | Prof. Mark Nitz |
| 127-IdU | S-Phase | - | Fluidigm |
| 128-Te | TOB*is* Barcode | - | Prof. Mark Nitz |
| 130-Te | TOB*is* Barcode | - | Prof. Mark Nitz |
| 141-Pr | Phospho-PDPK1 [S241] | J66-653.44.22 | BD Biosciences |
| 142-Nd | Cleaved-Caspase 3 [D175] | D3E9 | CST |
| 143-Nd | C-MYC | D84C12 | CST |
| 144-Nd | Phospho-MEK1/2 [S221] | 166F8 | CST |
| 145-Nd | Na/K-ATPase | EP1845Y | Abcam |
| 146-Nd | Phospho-MKK4/SEK1 [S257] | C36C11 | CST |
| 147-Sm | Phospho-BTK [Y551] | 24a/BTK | BD Biosciences |
| 148-Nd | Phospho-SRC [Y418] | SC1T2M3 | BD Biosciences |
| 149-Sm | Phospho-4E-BP1 [T37/46] | 236B4 | CST |
| 150-Nd | Phospho-RB [S807/811] | J112-906 | BD Biosciences |
| 151-Eu | Phospho-PKCα [T497] | K14-984 | BD Biosciences |
| 152-Sm | Phospho-AKT [T308] | J1-223.371 | BD Biosciences |
| 153-Eu | Phospho-CREB [S133] | 87G3 | CST |
| 154-Sm | Phospho-SMAD1 [S463/465]<br>Phospho-SMAD5 [S463/465]<br>Phospho-SMAD9 [S465/467] | D5B10 | CST |
| 155-Gd | Phospho-AKT [S473] | D9E | CST |
| 156-Gd | Phospho-NF-κB p65 [S529] | K10-895.12.50 | BD Biosciences |
| 157-Gd | Phospho-MKK3 [S189] / MKK6 [S207] | D8E9 | CST |
| 158-Gd | Phospho-p38 MAPK [T180/Y182] | D3F9 | CST |
| 159-Tb | Phospho-MAPKAPK2 [T334] | 27B7 | CST |
| 160-Gd | Phospho-AMPKα [T172] | 40H9 | CST |
| 161-Dy | Phospho-BAD [S112] | 40A9 | CST |
| 162-Dy | LRIG1 | Polyclonal | R&D Systems |
| 163-Dy | Phospho-p90RSK [T359] | D1E9 | CST |
| 164-Dy | Phospho- p120-Catenin [T310] | 22/p120 (pT310) | BD Biosciences |
| 165-Ho | β-Catenin [Active] | D13A1 | CST |
| 166-Er | Phospho-GSK-3β [S9] | D85E12 | CST |
| 167-Er | Phospho-ERK1/2 [T202/Y204] | 20A | BD Biosciences |
| 168-Er | Phospho-SMAD2 [S465/467]<br>Phospho-SMAD3 [S423/425] | D27F4 | CST |
| 169-Tm | Phospho-STAT3 [Y705] | 4/P-STAT3 | BD Biosciences |
| 170-Er | Arginase-1 | D4E3MTM | CST |
| 171-Yb | F4/80 | BM8 | BioLegend |
| 172-Yb | Phospho-S6 [S235/236] | D57.2.2E | CST |
| 173-Yb | Podoplanin (PDPN) | 8.1.1 | BioLegend |
| 174-Yb | RFP | 8E5.G7 | Rockland |
| 175-Lu | CD44 | IM7 | BioLegend |
| 176-Yb | Cyclin B1 | GNS-11 | BD Biosciences |
| 191-Ir | DNA | - | Fluidigm |
| 193-Ir | DNA | - | Fluidigm |
| 194-Pt | Dead Cells | - | Fluidigm |
| 196-Pt | TOB*is* Barcode | - | Fluidigm |
| 198-Pt | TOB*is* Barcode | - | Fluidigm |
| 209-Bi | CD68 | FA-11 | BioLegend |

Extracellular
Intracellular
Non-Antibody Parameter

#### Supplementary Table 3 – TOB*is* Matrix for Organoid Time-Course (Figure 2)

| | TeMal | | | | Cisplatin | |
|---|---|---|---|---|---|---|
| | $^{124}$Te | $^{126}$Te | $^{128}$Te | $^{130}$Te | $^{196}$Pt | $^{198}$Pt |
| **Day 1** | + | − | + | − | − | + |
| **Day 2** | + | − | − | + | − | + |
| **Day 3** | + | − | − | − | + | + |
| **Day 4** | − | + | + | + | − | − |
| **Day 5** | − | + | − | − | + | + |
| **Day 6** | − | − | + | − | + | + |
| **Day 7** | − | − | − | + | + | + |

#### Supplementary Table 4 – TOB*is* Matrix for CRC Tumour Microenvironment Model (Figure 3)

| Genotype | Microenvironment | TeMal | | | | Cisplatin | |
|---|---|---|---|---|---|---|---|
| | | $^{124}$Te | $^{126}$Te | $^{128}$Te | $^{130}$Te | $^{196}$Pt | $^{198}$Pt |
| **WT** | **Organoid** | + | − | − | − | + | + |
| | **Organoid / Macrophage** | + | + | − | − | − | + |
| | **Organoid / Fibroblast** | − | − | + | + | + | − |
| | **Organoid / Fibroblast / Macrophage** | + | − | + | + | − | − |
| **A** | **Organoid** | + | − | + | − | + | − |
| | **Organoid / Macrophage** | + | − | − | + | + | − |
| | **Organoid / Fibroblast** | + | − | + | − | − | + |
| | **Organoid / Fibroblast / Macrophage** | + | − | − | + | − | + |
| **AK** | **Organoid** | + | + | + | − | − | − |
| | **Organoid / Macrophage** | − | + | + | − | + | − |
| | **Organoid / Fibroblast** | − | + | + | + | − | − |
| | **Organoid / Fibroblast / Macrophage** | − | + | + | − | − | + |
| **AKP** | **Organoid** | − | + | − | + | + | − |
| | **Organoid / Macrophage** | − | + | − | − | + | + |
| | **Organoid / Fibroblast** | − | + | − | + | − | + |
| | **Organoid / Fibroblast / Macrophage** | + | + | − | − | + | − |
| | **Macrophage** | − | − | + | − | + | + |
| | **Fibroblast** | − | − | + | + | − | + |
| | **Fibroblast / Macrophage** | − | − | − | + | + | + |

#### SUPPLEMENTARY FIGURE LEGENDS

**Supplementary Figure 1 – Rare-Earth Metal-Conjugated Antibodies for Organoid Cell-Type Identification.**

**a)** Summary of reagents used in organoid directed differentiation.

**b)** Fuzzy logic diagram of organoid directed differentiation.

**c)** Confocal immunofluorescence (IF) of small intestinal organoids stained with rare-earth metal-conjugated mass cytometry (MC) antibodies showing individual cell-type markers (red), F-Actin (white), and DAPI (blue) following directed differentiation, scale bars = 50 µm.

**d)** UMAP (Uniform Manifold Approximation and Projection) distributions of 20,000 single organoid cells analysed by MC following directed differentiation, demonstrating the specificity of rare-earth metal-conjugated antibodies for organoid cell-type identification.

**e)** Force-directed Scaffold maps constructed with cell-type landmarks identified from untreated small intestinal organoids. Unsupervised distribution of single organoid cells following directed differentiation demonstrates the specificity of rare-earth metal-conjugated antibodies for organoid cell-type identification.

**f)** Earth Mover's Distance (EMD) quantification of cell-type identification markers from MC analysis of *Lgr5-EGFP-ires-CreERT2* small intestinal crypt-cells. LGR5-GFP$^+$ stem cells are positive for LRIG1 whereas negative for all differentiated cell-type markers. LRIG1$^+$ cells are LGR5-GFP-positive, but negative for all differentiated cell-type markers.

**Supplementary Figure 2 – Cell-Type and Cell-State Identification.**

**a)** Cell-type identification. Raw MC data is gated for Gaussian parameters, DNA$^{high}$, Cisplatin$^{low}$, and EpCAM$^{+}$ / Pan-CK$^{+}$ to identify single epithelial cells. Cells are then gated to remove doublets stained positive for mutually exclusive cell-type identification and cell-state markers. Single cells are clustered and visualised in UMAP space and each cell-type is gated by their identification markers.

**b)** Cell-state classification. Single cells are resolved into cell-states including apoptotic (pRB$^{-}$, cC3$^{+}$), G0- (pRB$^{-}$, cC3$^{-}$), S- (pRB$^{+}$, IdU$^{+}$), M- (pRB$^{+}$, IdU$^{-}$, pHH3$^{+}$), G2- (pRB$^{+}$, IdU$^{-}$, pHH3$^{-}$, Cyclin B1$^{+}$), and G1-phase (pRB$^{+}$, IdU$^{-}$, pHH3$^{-}$ , Cyclin B1$^{-}$).

**c)** EMD quantification of cell-type identification markers across all small intestinal organoid cell-types from Figs. 1 and 2. Each epithelial cell-type is enriched for its respective cell-type identification marker.

**d)** Cell-state quantification of all small intestinal organoid cell-types from Figs. 1 and 2. Stem, Paneth, and enteroendocrine cells are generally proliferative, whereas tuft, goblet cells, and enterocytes are often quiescent or apoptotic.

**Supplementary Figure 3 – Cell-Type and Cell-State Specific Signalling Networks in Intestinal Organoids.**

Cell-type and cell-state specific signalling networks of 27 PTMs from 1 million single organoid cells analysed by MC, with nodes coloured by PTM-EMD scores quantifying PTM intensity (relative to all organoid cells), and edges coloured by DREMI scores quantifying PTM-PTM connectivity.

**Supplementary Figure 4 – Maxpar versus TOB*is* Organoid Barcoding *in situ.***

**a)** De-barcoded cell counts from small intestinal organoids barcoded using the Maxpar Cell-ID™ Barcoding Kit either after removal from Matrigel (*ex situ*) or while still in Matrigel (*in situ*). The Maxpar kit can only be used to barcode organoid cells *ex situ* (two-tailed unpaired *t*-test, $p = 0.0002$). Error bars represent standard deviation (SD) of two technical replicates.

**b)** Mean dual counts of $^{110}$Pd from Matrigel incubated in PBS, Maxpar Cell-ID™ barcode #20 ($^{106}$Pd, $^{108}$Pd, and $^{110}$Pd) resuspended in PBS (PBS + BC), or Maxpar Cell-ID™ barcode #20 resuspended in Maxpar Cell Staining Buffer (CSB + BC). While Maxpar barcodes react with Matrigel, the reaction is fully blocked by the presence of CSB (two-tailed unpaired *t*-test, $p < 0.0001$). Error bars represent SD of three technical replicates.

**c)** Maximum-gain confocal IF images of NHS ester- / $C_2$ maleimide-Alexa 647 fluorescent probes shown in Fig. 3a to highlight Matrigel staining. High Matrigel background staining is observed with amine-reactive probes, whereas thiol-reactive probes only bind organoids.

**d)** Small intestinal organoids concurrently stained *in situ* with Maxpar Cell-ID™ ($^{102}$Pd, $^{104}$Pd, $^{105}$Pd, $^{106}$Pd, $^{108}$Pd, and $^{110}$Pd) and TOB*is* 1—20 ($^{124}$Te, $^{126}$Te, $^{128}$Te, $^{130}$Te, $^{196}$Pt, and $^{198}$Pt) barcodes using analogous 20-plex *6*-choose-*3* matrices. While Maxpar barcodes are unsuitable for labelling organoids in Matrigel, TOB*is* barcodes capably resolve each organoid condition.

**Supplementary Figure 5 – Maxpar versus TOB*is* Cell-Recovery Comparison.**

**a)** Example 10-plex barcoding workflow for organoid MC using Maxpar Cell-ID™ Barcoding Kit. As it requires organoids to be individually removed from Matrigel, individually dissociated, permeabilised, and quenched, at least 7 centrifugation steps are required per *n* organoid sample(s) (total = 7*n*). This increases the chance of cell-loss and is compounded as the sample *n* increases.

**b)** Example 10-plex barcoding workflow for organoid MC using TOB*is*. As TOB*is* barcodes are applied and quenched *in situ*, no centrifugation or permeabilisation steps are required for barcoding. This increases sample-throughput and reduces cell-loss. Following a pooled dissociation, two blocking / centrifugation steps are performed prior to rare-earth metal antibody staining irrespective of sample *n*.

**c)** and **d)** Dissociation and centrifugation steps (*s*) required for multiplexed organoid experiments using Maxpar or TOB*is* barcoding illustrating the impracticability of *ex situ* barcoding for large-scale organoid experiments (e.g. screening applications).

**e)** Single-cell recovery of serially-titrated small intestinal organoids barcoded with either Maxpar (*ex situ*) or TOB*is* (*in situ*). By reducing the number of dissociation and centrifugation steps, 4.8-fold more single cells were recovered by TOB*is* compared to Maxpar barcoding (two-tailed ratio-paired *t*-test, $p < 0.001$). Error bars represent standard error of the mean (SEM) of two technical replicates.

**f)** Cell-type recovery (cell count) from equally seeded small intestinal organoids analysed by MC and de-barcoded by Maxpar or TOB*is*. Cell-types were manually identified in UMAP space. Error bars represent SD of two technical replicates.

**g)** Cell-type recovery (population percentage) from equally sampled small intestinal organoid cells analysed by MC and de-barcoded by Maxpar or TOB*is* (linear regression of correlation, $R^2 = 0.86$). Cell-types were manually identified in UMAP space. Error bars represent SD of two technical replicates.

**h)** Mean debarcoded cell counts in the absence or presence of glutathione (GSH) in PBS washes following TOB*is* barcoding. GSH *in situ* washes quench unbound barcodes before pooling and increase TOB*is* debarcoding efficiency (two-tailed unpaired t-test, $p < 0.0001$). Error bars represent SD of three technical replicates.

**Supplementary Figure 6 – Heterocellular Colorectal Cancer (CRC) Organoid Cells.**

**a)** Confocal IF of CRC organoid genotypes (wild-type (WT), *shApc* (A), *shApc* and *Kras*$^{G12D/+}$ (AK), and *shApc*, *Kras*$^{G12D/+}$ and *Trp53*$^{R172H/-}$ (AKP)) stained with Pan-CK, LRIG1, CD44, and C-MYC (red), F-Actin (white), and DAPI (blue), scale bars = 50 μm. Pan-CK and LRIG1 (in combination with CEACAM1) were subsequently used for epithelial / stem cell identification in MC.

**b)** Confocal IF of colonic fibroblasts cultured in 2D (plastic) and 3D (Matrigel) demonstrates positive staining for fibroblast markers PDPN, Vimentin, PDGFRα, FOXL1, and GLI1 (red), F-Actin (white), and DAPI (blue), scale bars = 50 μm. PDPN and RFP were subsequently used for fibroblast identification in MC.

**c)** Confocal IF of primary macrophages cultured in 2D (plastic) and 3D (Matrigel) demonstrates positive staining for macrophages markers CD68, F4/80, CD45, CD11b, and CX3CR1 (red), F-Actin (white), and DAPI (blue), scale bars = 50 μm. CD68 and F40/80 were subsequently used for macrophage identification in MC.

**Supplementary Figure 7 – CRC Organoid Signalling Networks in the CRC Tumour Microenvironment (TME) Model.**

Signalling networks of 28 PTMs in colonic organoid cells from each genotype (wild-type (WT), *shApc* (A), *shApc* and *Kras*$^{G12D/+}$ (AK), or *shApc*, *Kras*$^{G12D/+}$, and *Trp53*$^{R172H/-}$ (AKP)) and microenvironment condition in the CRC TME model analysed by MC. Signalling nodes are connected by DREMI edges and PTM activity is displayed as EMD relative to all cells across all conditions.

**Supplementary Figure 8 – Macrophage and Fibroblast Signalling Profiles in the CRC TME Model.**

**a)** and **b)** Signalling networks of 28 PTMs in macrophages and colonic fibroblasts from each genotype and microenvironment condition in the CRC TME model analysed by MC. Signalling nodes are connected by DREMI edges and PTM activity is displayed as EMD relative to all cells across all conditions.

**c)** and **d)** PCA of 28 PTM-EMDs for macrophages and colonic fibroblasts across all genotype / microenvironment combinations.

**e)** and **f)** PCA of 756 PTM-DREMIs for macrophages and colonic fibroblasts across all genotype / microenvironment combinations.

**Supplementary Figure 9 – CRC Organoid Microenvironment and Genotype Scaffold Maps.**

**a)** Force-directed Scaffold maps were constructed from WT colonic organoids alone or co-cultured with macrophages and/or colonic fibroblasts. Unsupervised distribution of WT, A, AK, and AKP colonic organoid monocultures and co-cultures with macrophages and/or colonic fibroblasts revealed that cell-autonomous oncogenic mutations mimic microenvironment-driven signalling.

**b)** Force-directed Scaffold maps were constructed from WT, A, AK, and AKP colonic organoid monocultures. Unsupervised distribution of WT, A, AK, and AKP colonic organoid monocultures and co-cultures with macrophages and/or colonic fibroblasts confirmed that oncogenic signalling profiles mimic microenvironment-driven signalling.

### Supplementary Figure 1

**a**

| Abbreviation | Ligand | Signalling | Target |
|---|---|---|---|
| E | EGF | MAPK ON | EGFR |
| N | Noggin | BMP OFF | BMP |
| R | R-Spondin-1 | WNT ON | LGR5/ZNRF |

| Abbreviation | Inhibitor | Signalling | Target |
|---|---|---|---|
| C | CHIR99021 | WNT ON | GSK-3β |
| I | IWP-2 | WNT OFF | PORCN |
| V | Valproic Acid | Notch ON | HDAC |
| D | DAPT | Notch OFF | γ-Secretase |

**b**

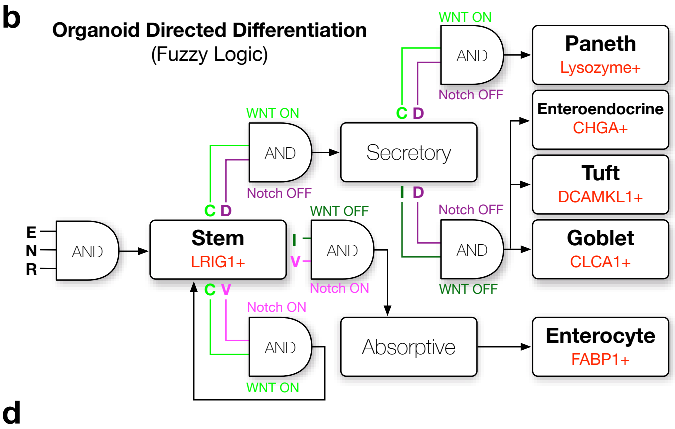


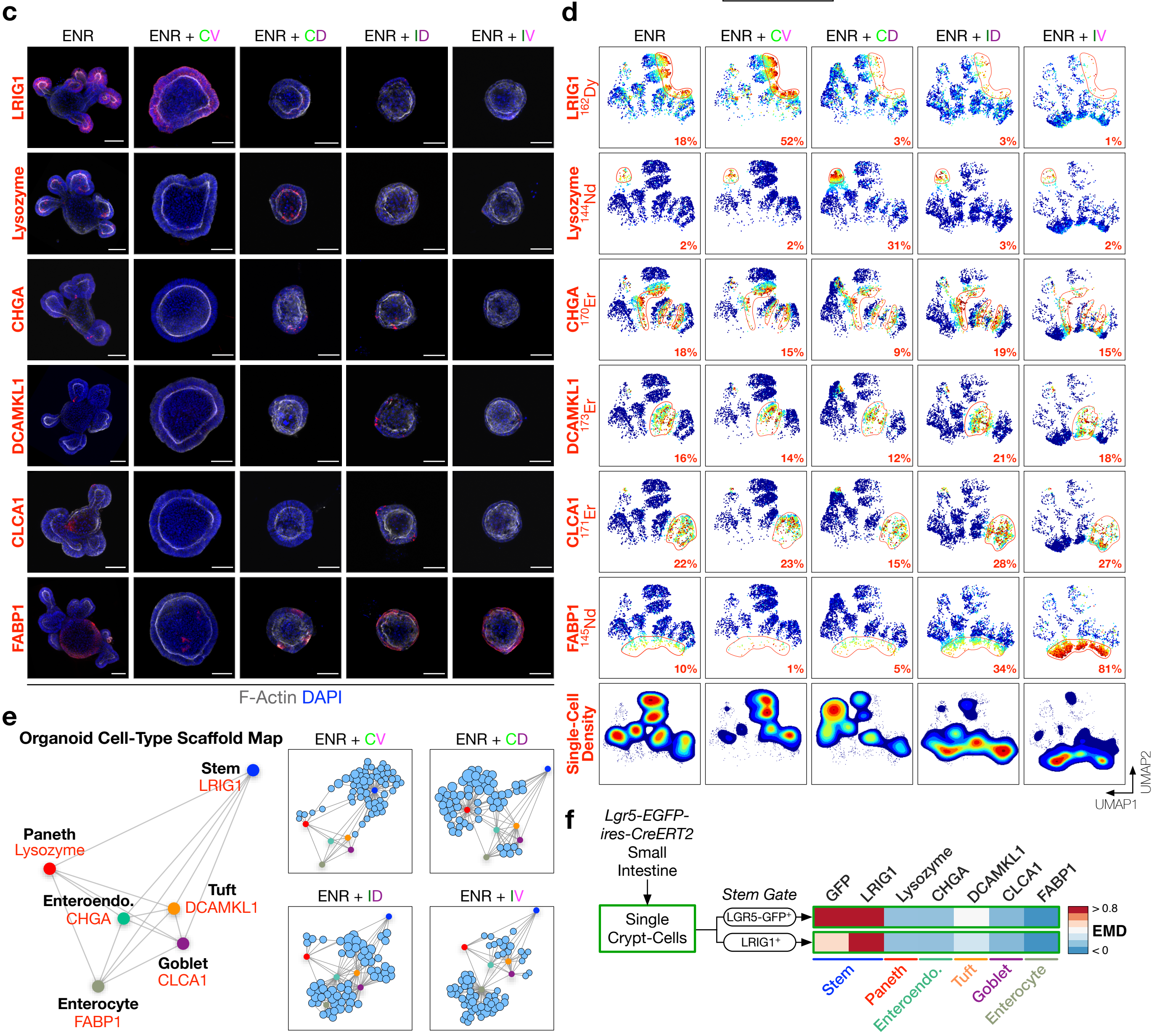

### Supplementary Figure 2

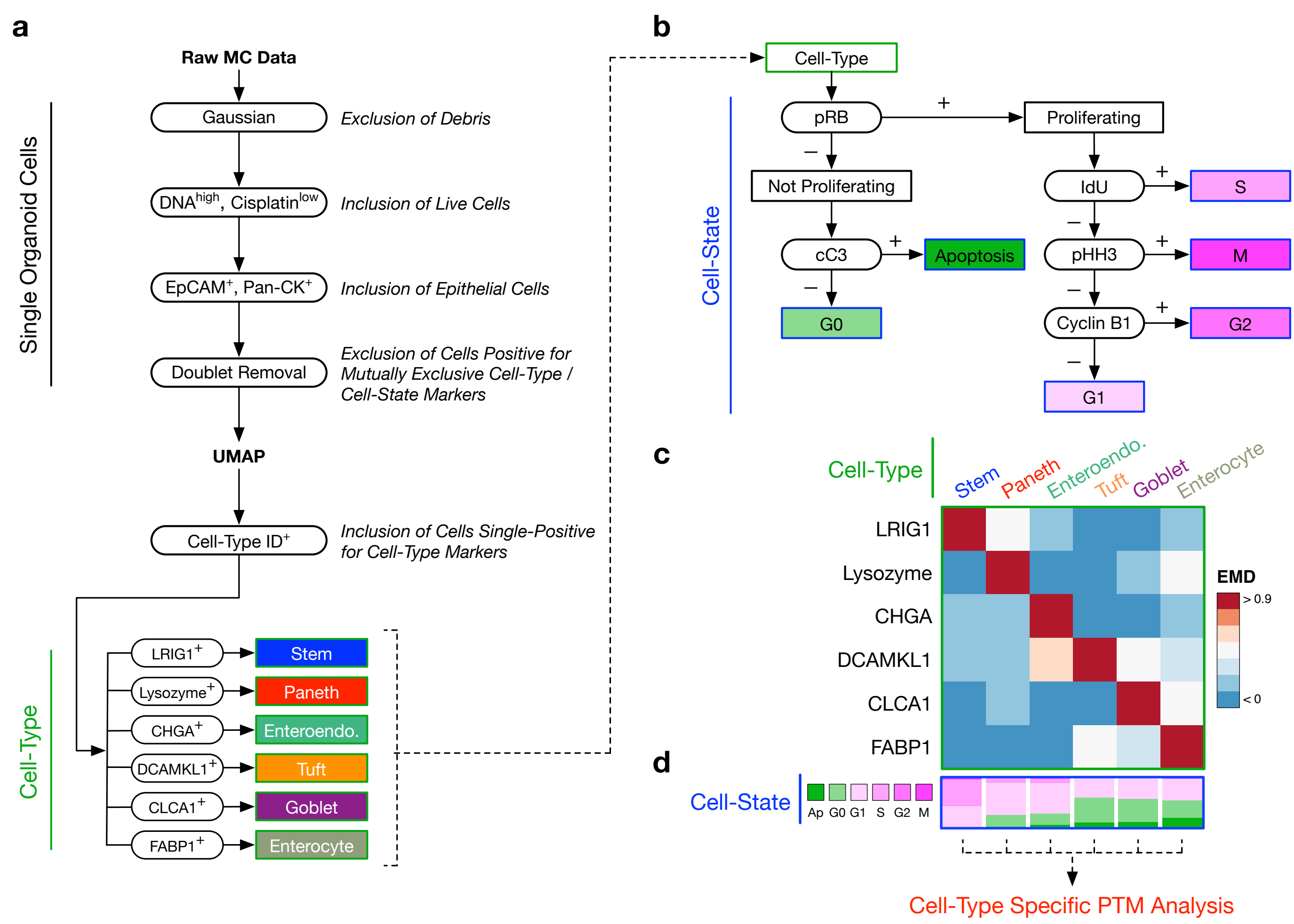

### Supplementary Figure 3

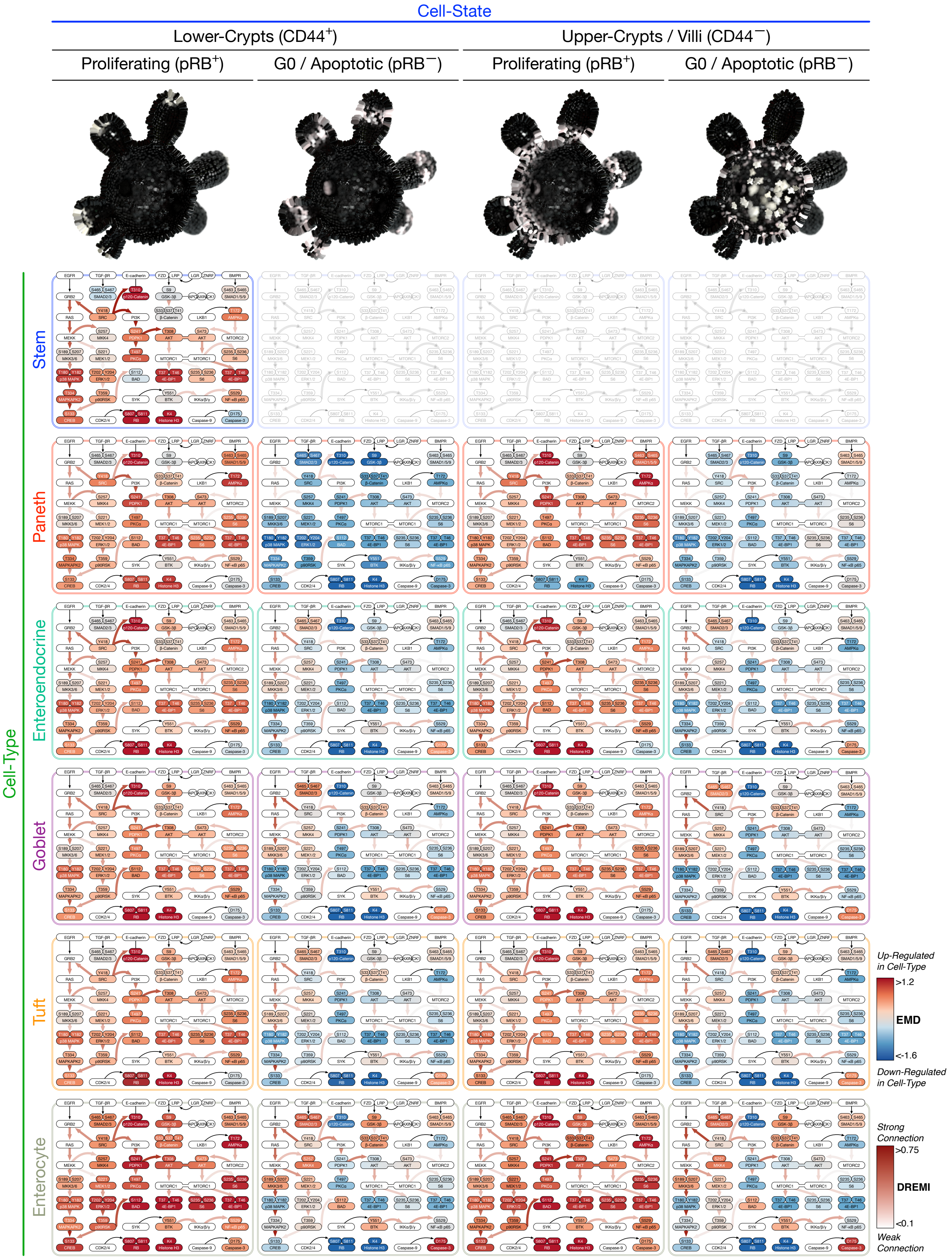

### Supplementary Figure 4

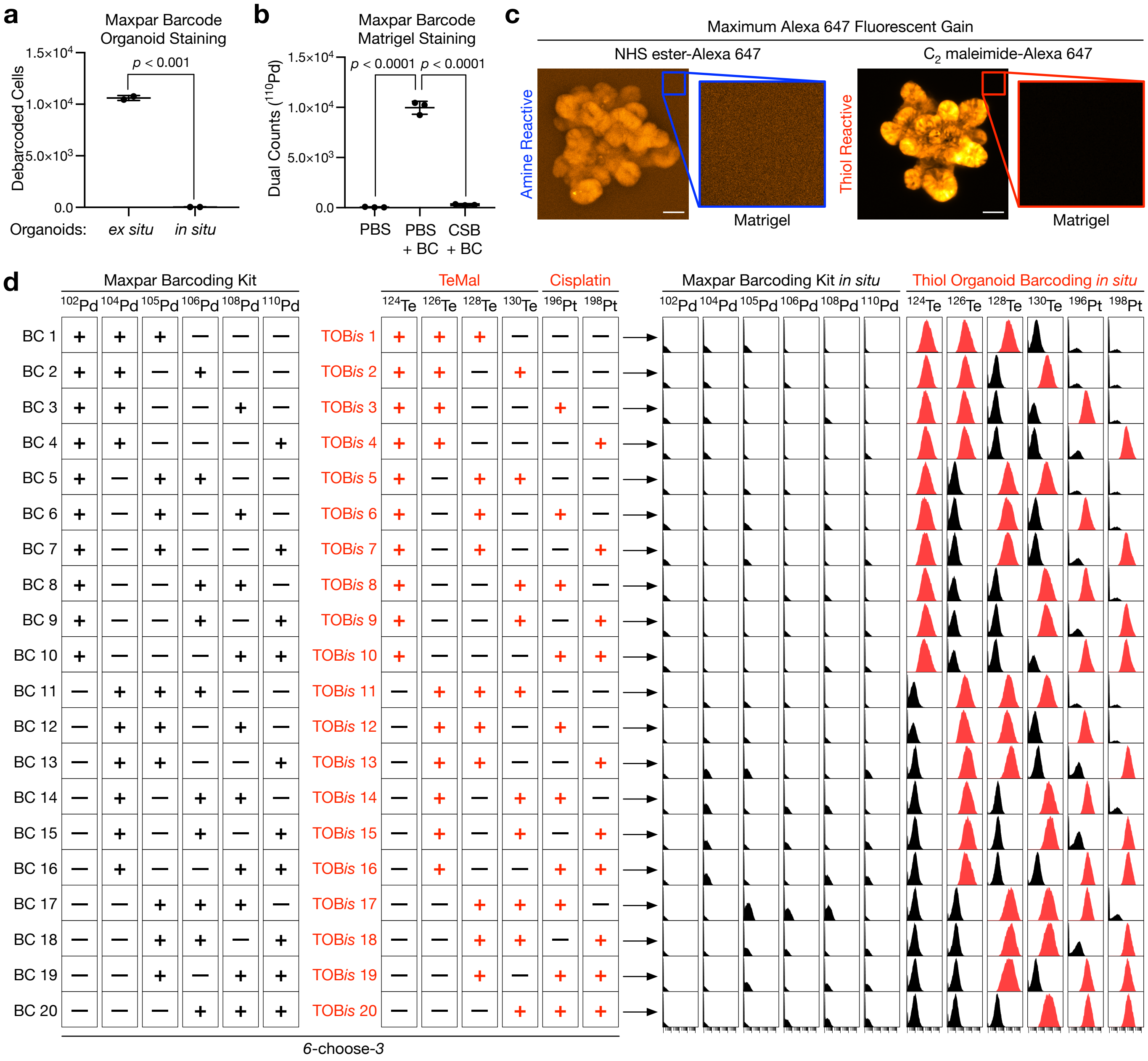

### Supplementary Figure 5

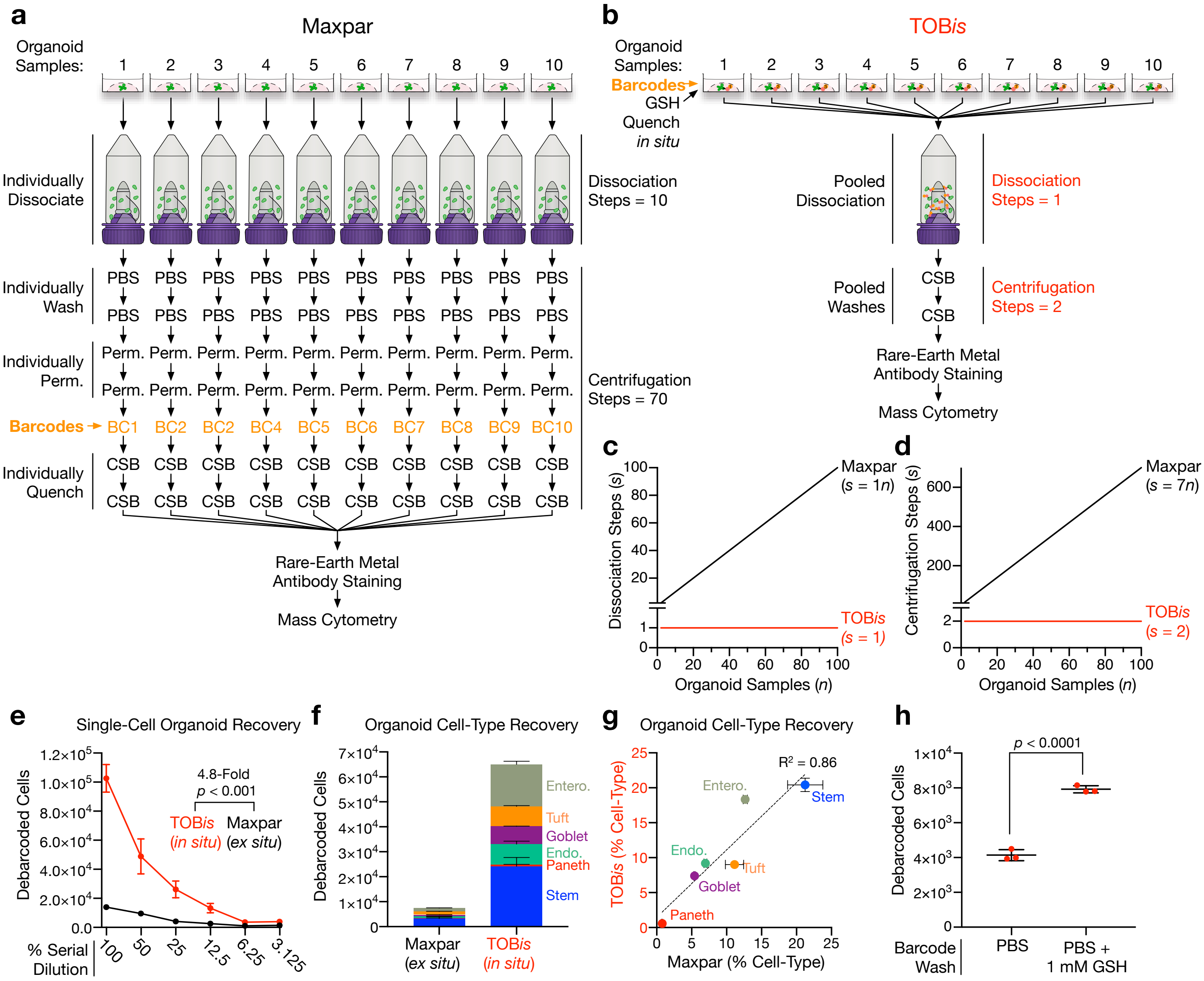

### Supplementary Figure 6

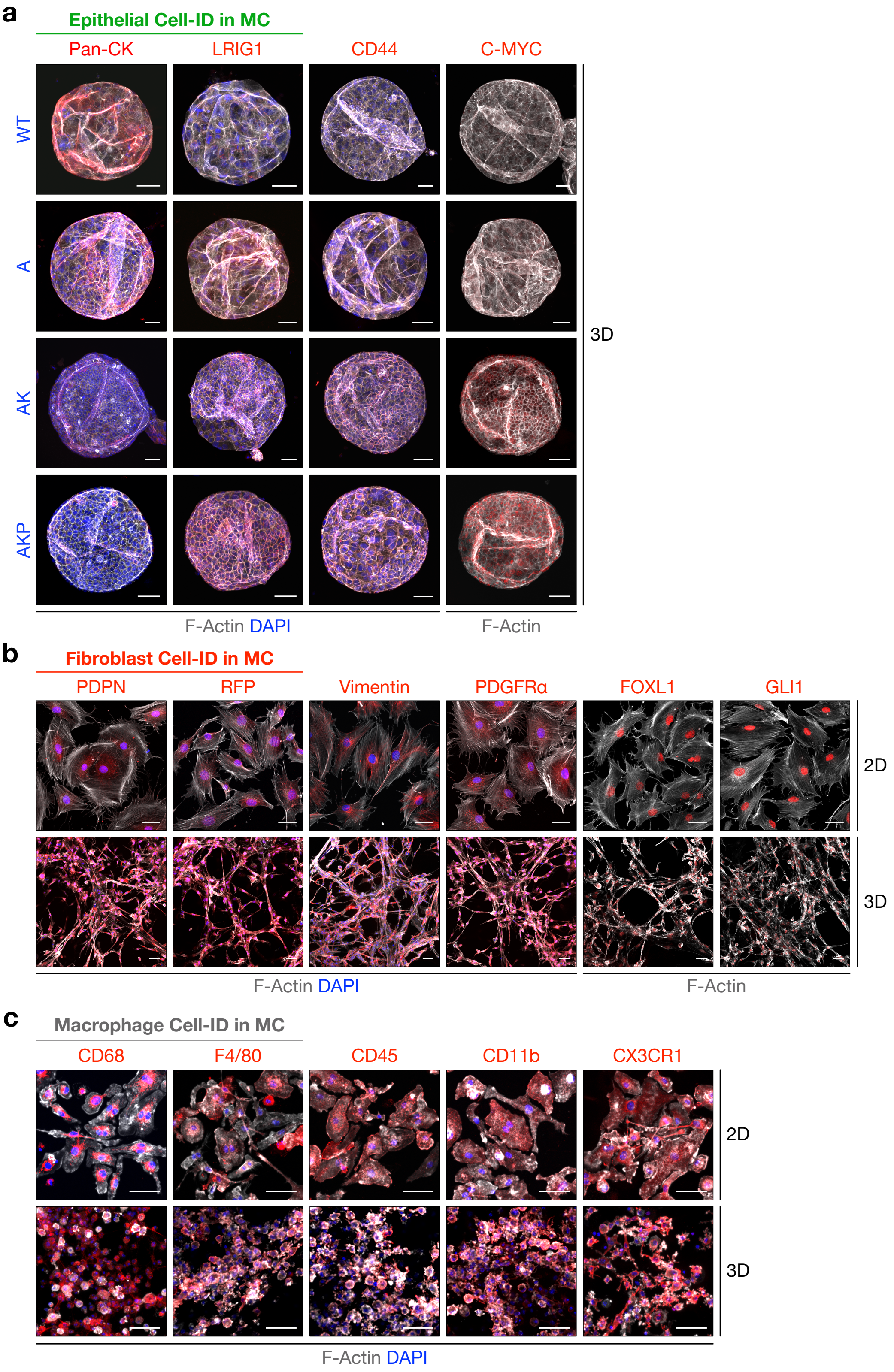

### Supplementary Figure 7

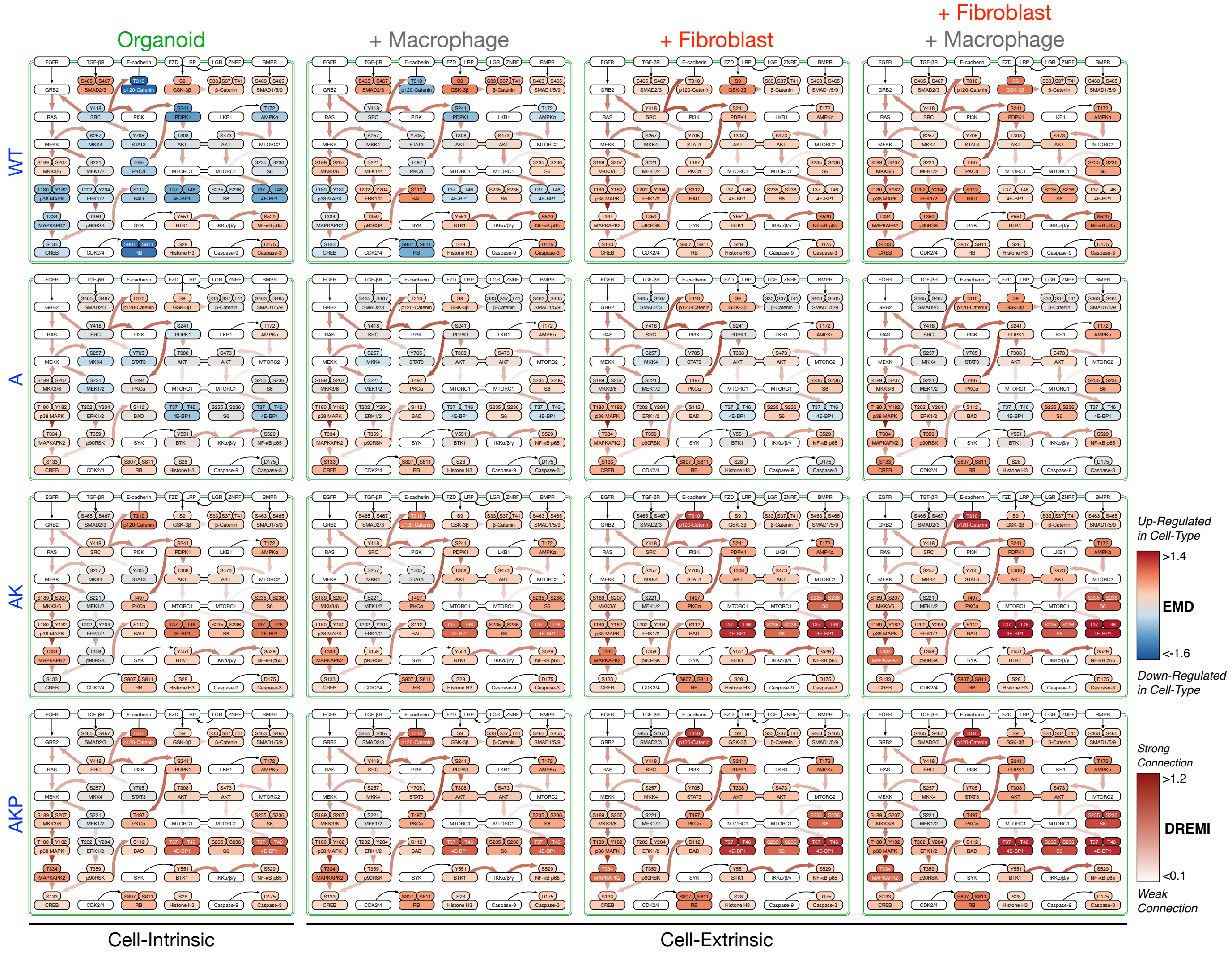

### Supplementary Figure 8

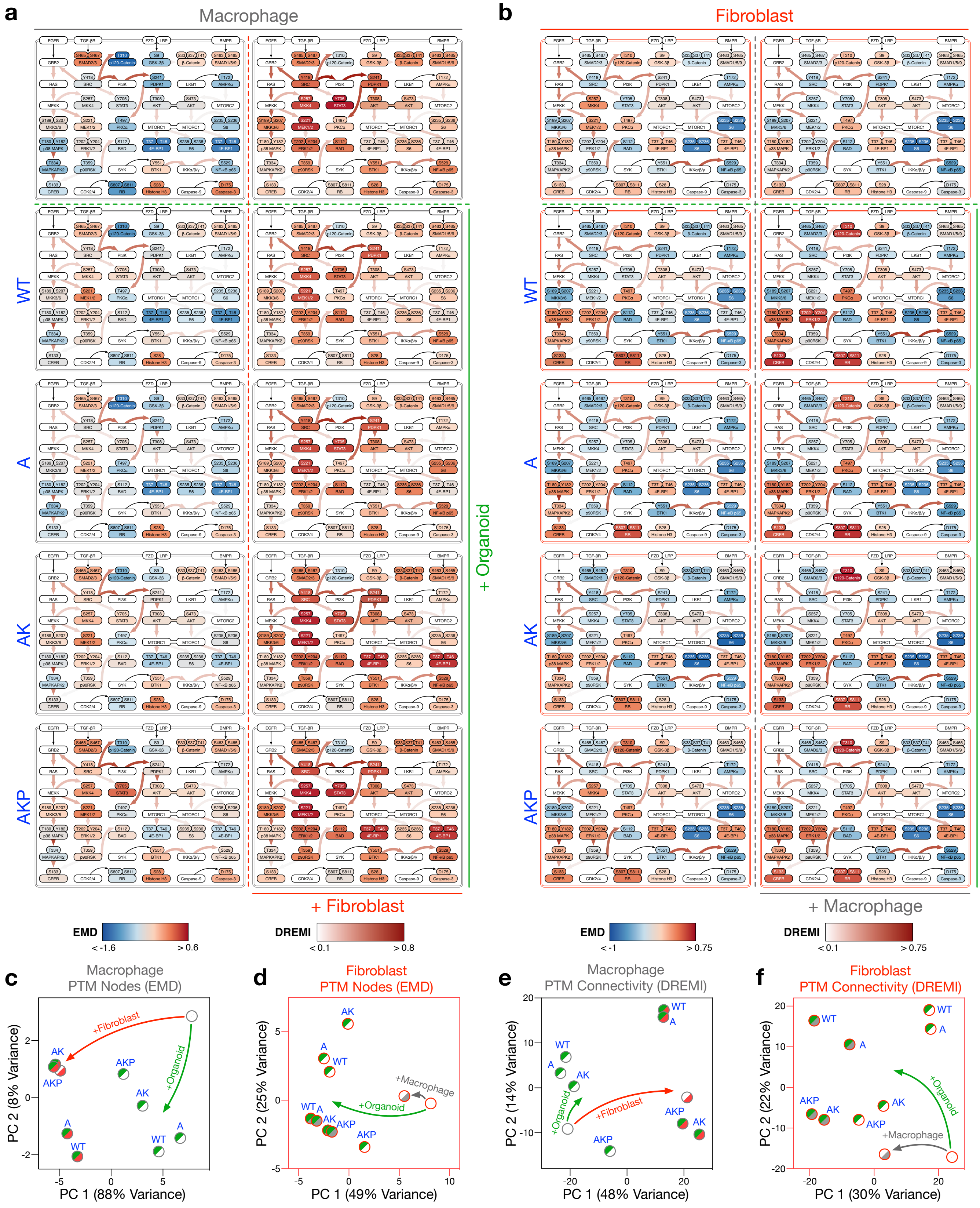

### Supplementary Figure 9

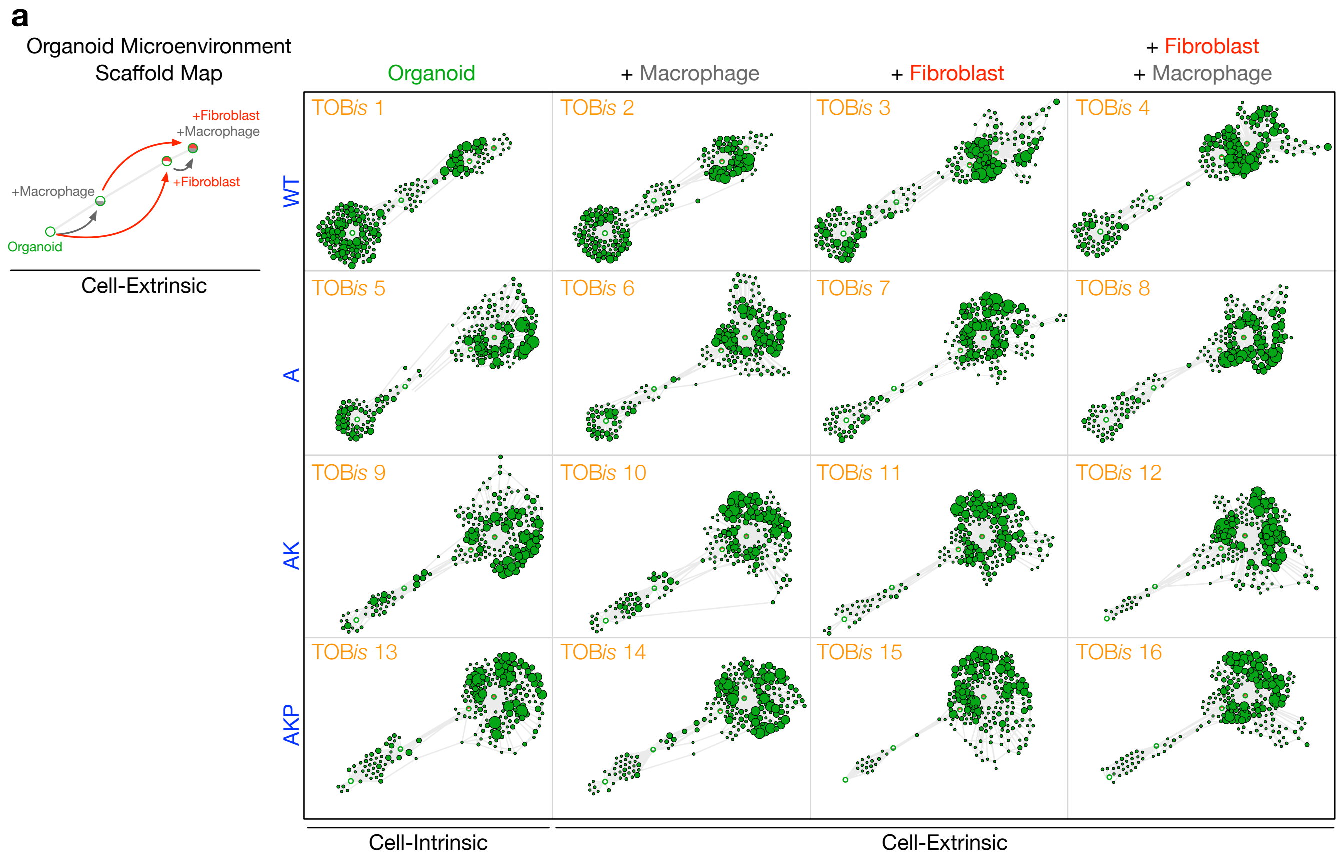


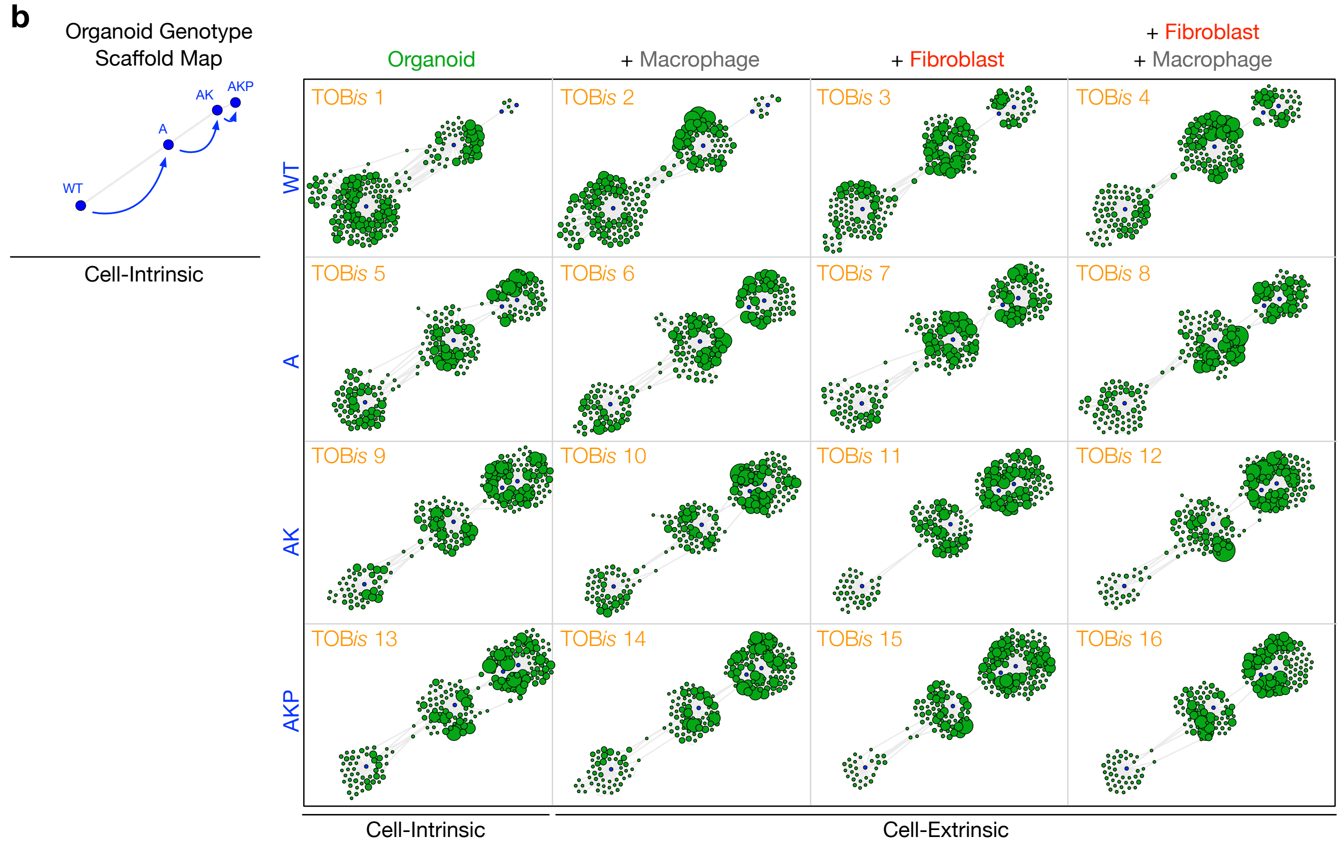
